# Structural and mechanistic insights into SLC34 phosphate import

**DOI:** 10.64898/2026.05.25.727688

**Authors:** Qinyu Zhu, Omar Almakki, Melinda M. Diver

## Abstract

Dysregulation of inorganic phosphate (Pi) homeostasis contributes to metabolic disease, cancer, pathological calcification, and kidney disease. Systemic phosphate balance is regulated by SLC34 transporters that mediate renal Pi retention (SLC34A1/A3) and intestinal dietary Pi absorption (SLC34A2). SLC34s couple Pi uptake to the symport of sodium (Na^+^) down its electrochemical gradient. Mutations or altered expression of SLC34 proteins are linked to disorders such as chronic kidney disease (CKD), where hyperphosphatemia is a major complication, and the lung disease pulmonary alveolar microlithiasis (PAM), caused by inactivating SLC34A2 mutations. SLC34A2 is also overexpressed in most ovarian and uterine tumors, making it an attractive target for antibody-drug conjugates. We present cryo-EM structures of SLC34A2 when the transporter is empty, bound to Na^+^ ions only, fully loaded with Na^+^ ions and Pi, and bound to an inhibitor phosphonoformic acid (PFA), revealing its distinct architecture, substrate and ion binding sites, the role of Na^+^, and multiple transporter states. Pi binds at a highly symmetric, membrane-embedded pocket positioned approximately mid-membrane and is coordinated by its signature QSSS repeat motifs. Na^+^ shapes the Pi-binding pocket and drives the transition from the outward-open to occluded state. Integrated with functional analyses, these structures reveal that SLC34 transporters operate through an atypical alternating access cycle defined by coordinated elevator movements of an auxiliary gate domain. This work lays a foundational framework for understanding Pi regulation and opens new avenues for therapeutic strategies targeting disorders linked to phosphate imbalance.

## INTRODUCTION

Maintaining inorganic phosphate (Pi) homeostasis is essential for numerous physiological processes, including energy metabolism, biomolecule synthesis, and skeletal mineralization, and is therefore critical for overall human health. The kidneys and small intestine act in concert to tightly regulate circulating levels of Pi by balancing intestinal absorption, skeletal storage, and kidney retention or excretion^1^. These actions are accomplished by sodium (Na^+^)-dependent Pi importers belonging to the SLC34 solute carrier family. SLC34A1 and A3 are responsible for Pi reabsorption in the renal proximal tube^2,3^.

SLC34A2, also known as NaPi-IIb, plays a crucial role in Pi absorption in the intestines by facilitating the uptake of Pi from the digestive tract^4-8^. SLC34A2 is also more broadly expressed, including in the lungs and secretory glands such as salivary, thyroid, and mammary glands^9^, and consequently plays a wider role in physiological function and disease.

SLC34A2 is implicated in several human diseases. Chronic kidney disease (CKD) afflicts more than 800 million people worldwide and frequently progresses to late-stage renal failure and increased mortality^10^. A major complication of CKD is hyperphosphatemia, characterized by elevated serum phosphate levels that contribute to bone weakening, pathological calcium phosphate deposits in multiple organs, cardiovascular disease, and further kidney damage^11,12^. Consequently, there is strong clinical interest in targeting SLC34 transporters to reduce systemic Pi burden. Inactivating mutations in SLC34A2 cause pulmonary alveolar microlithiasis (PAM), a disease in which the formation of calcifications in the lung diminishes its function, and for which there are no effective treatments^13^. Furthermore, SLC34A2 is overexpressed in ∼80-90% of ovarian tumors^14^, and antibody-drug conjugates (ADCs) targeting SLC34A2 are currently being evaluated in clinical trials as cancer-selective delivery systems for cytotoxic therapeutics^15^. Ovarian cancer is often diagnosed at a late stage, making it challenging to treat, and recurrence remains common. The first-line therapy for ovarian cancer is typically a combination of surgery and chemotherapy, and there is an urgent clinical need to discover novel strategies for detection and targeted treatment.

Although SLC34 transporters are integral to human physiology, their structures have remained elusive. The SLC34 family lacks sequence conservation with other transporters, and computational predictions suggest a novel structural architecture. SLC34 transporters share an inverted repeat topology and a signature QSSS motif, which are thought to play a critical role in defining the transport pathway. However, despite extensive physiological characterization, the absence of experimentally resolved structures has limited our understanding of SLC34 domain organization and molecular transport mechanisms. Here, we close this gap by resolving a series of single-particle cryoelectron microscopy (cryo-EM) structures of SLC34A2 that unveil a distinct transporter architecture and, together with complementary functional analyses, reveal the molecular principles governing Pi and Na^+^ recognition, the role of Na^+^, and their coupling mechanisms. We capture both outward-open and occluded conformations, providing key snapshots along the alternating access transport cycle that support an unexpected mechanism in which a peripheral gate domain, not the transport domain, serves as the primary mobile element for this atypical elevator transporter. This work provides a critical molecular framework for understanding relevant disease mechanisms and developing targeted therapeutic strategies.

## RESULTS

### Architecture of SLC34A2

We found that SLC34A2a from zebrafish (zfSLC34A2) exhibits superior conformational homogeneity in detergent-containing solutions in comparison to other orthologs. Human and zebrafish SLC34A2 share substantial sequence conservation (70% amino acid similarity) (Fig. S1), and the protein function of the zfSLC34A2 isoforms has been well characterized using transport assays and electrophysiology^16-18^, indicating that they exhibit key functional properties of the SLC34 family. Full-length zfSLC34A2 protein was engineered with an epitope tag inserted into the TM3-TM4 extracellular loop to facilitate binding of the ovarian-cancer-directed antibody MX35 (referred to here as zfSLC34A2-MX35 or SLC34A2), and this construct was used throughout our structural analyses (Fig. S2A). This SLC34A2 construct exhibits robust import of Pi when heterologously expressed in *Xenopus* oocytes, comparable to that of the wild-type protein (Fig. 1D and Fig. S3). Cryo-EM analysis of SLC34A2 alone produced only low-resolution maps (around 6 Å) likely owing to its small 70 kDa molecular weight and the absence of structured extramembrane domains that could help to resolve signal ambiguities. We first attempted to use the therapeutic MX35 Fab as a fiducial, however its binding epitope resides in a conformationally flexible loop and therefore does not adopt a fixed spatial orientation relative to SLC34A2 (Fig. S2B). Therefore, we generated monoclonal antibodies as fiducial markers. With the assistance of one of these Fabs, 12H07, we determined multiple single particle cryo-EM maps of SLC34A2 including under apo, Na^+^-bound, and Pi- and Na^+^-bound conditions with average resolutions up to 3.0 Å (Tab. 1 and Figs. S4-11). The cryo-EM maps reveal substantial conformational heterogeneity, with the highest resolution (∼2.5 Å) achieved in the core transport domain where substrate and ions bind (Fig. S6H).

**Figure 1.**
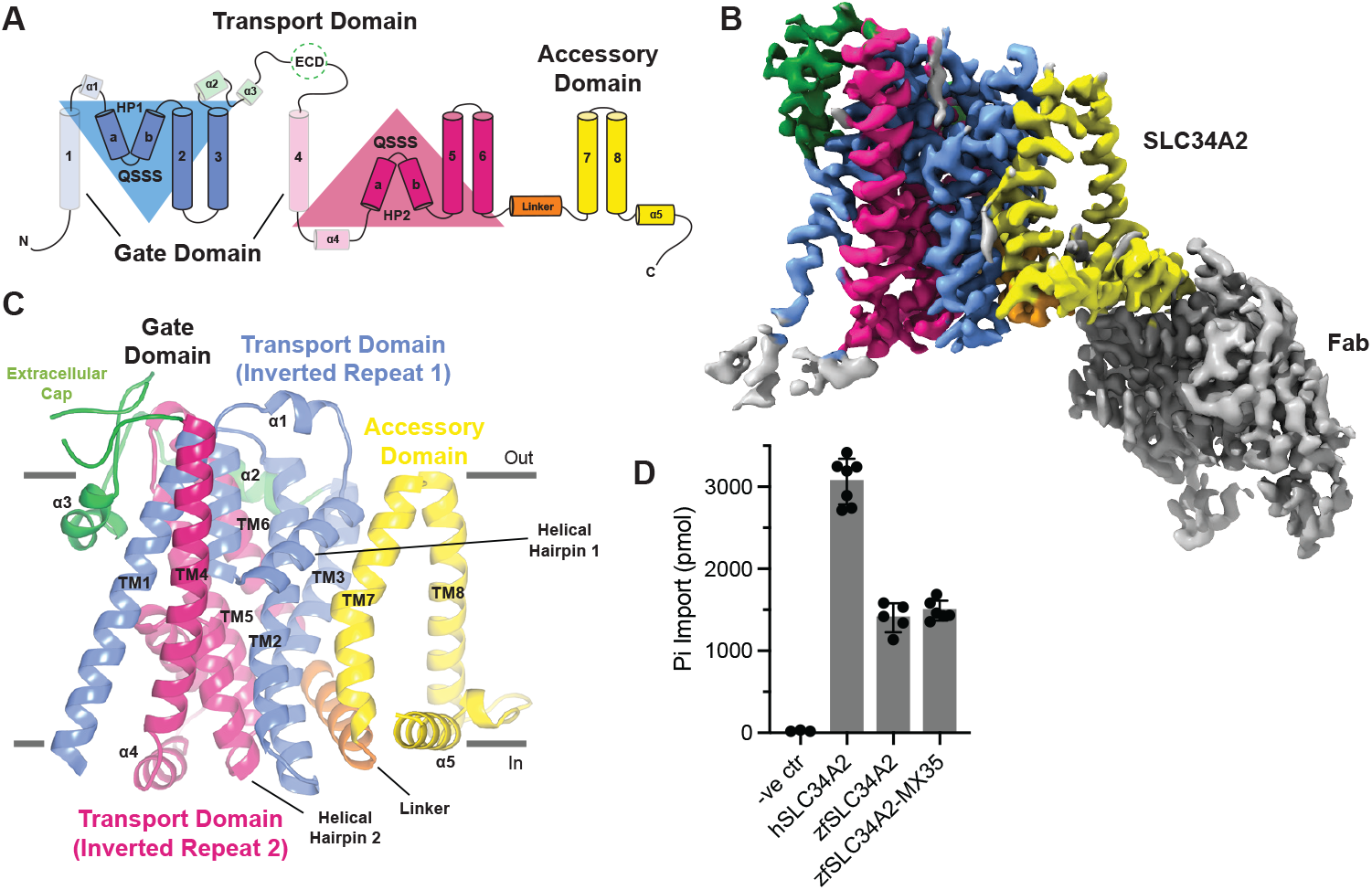
Architecture of SLC34A2. (A) Topology of SLC34A2. TM1, HP1, TM2-3 and TM4, HP2, TM5-6 form a pseudo-twofold symmetry, as denoted by the triangles. The gate domain is transparent. Inverted repeat domain 1 is colored blue, inverted repeat domain 2 is colored pink, the extracellular domain is colored green, the linker helix is colored orange, and the accessory domain is colored yellow. ECD, extracellular domain. (B) Cryo-EM density map of SLC34A2 colored identically to A. The Fab is colored gray. (C) Cartoon representation of the SLC34A2 structure, viewed parallel to the membrane plane. (D) Pi import mediated by the human and zebrafish orthologs of SLC34A2 following their overexpression in oocytes. zfSLC34A2-MX35 contains the binding epitope for the therapeutic MX35 SLC34A2 antibody. In oocytes expressing SLC34A2, application of 1 mM Pi, in the presence of 96 mM Na^+^, evoked radioactive ^32^P import. The negative control (-ve ctr) is uninjected oocytes. Data are represented as the mean ± SD (n=3-7 samples).

The SLC34 family adopts a novel solute carrier family (SLC) fold (Fig. 1A-C). A Dali search against experimental structures in the Protein Data Bank (PDB) and predicted structures in the AlphaFold Database yields no matches^19^. We resolved a monomeric structure, contrary to previous reports suggesting multimerization^20,21^, but consistent with the established functional unit of SLC34 family members^22^. SLC34A2 contains eight transmembrane α-helices (TM1-TM8) and two re-entrant helical hairpins (HP1 and HP2), which traverse halfway through the membrane before turning back. It features a structural repeat with an inverted topology, inclusive of TM1-HP1-TM2-TM3 (inverted repeat 1) and TM4-HP2-TM5-TM6 (inverted repeat 2). The structure is comprised of three main domains: central transport (TM2-3 and TM5-6, together with HP1 and HP2), peripheral gate (TM1 and TM4, alongside several extracellular α-helices (ECD) and an intracellular α-helix (α4)) and accessory (TM7-8). The extracellular loop recognized by MX35 binds is not visible. The Fab binds to the accessory domain, and we observed no effect on SLC34A2 function upon complex formation (Fig. 1B and Fig. S12).

### Phosphate-binding site

SLC34 transporters are known to translocate divalent Pi (HPO_4_^2-^)^23,24^, displaying Michaelian kinetics with respect to substrate, consistent with the translocation of one Pi per transport cycle^23,25^ and exhibit apparent substrate affinities of 100-350 µM^26-28^, suggesting a high affinity binding site. To determine where substrate binds and how substrate specificity and stoichiometry is maintained by SLC34A2, we determined structures of SLC34A2 in the presence of 5 mM Pi and 150 mM Na^+^. Na^+^ binding is a prerequisite of Pi binding^29^. Interestingly, in our structure, the two re-entrant helical hairpins, HP1 and HP2, that converge in the membrane core, enclose a non-protein density that we attribute to bound Pi substrate (Fig. 2A,C and Fig. S11A). The Pi binding site exhibits high symmetry that arises from the internal transporter repeats and is comprised of the signature QSSS repeat motifs (158-QSSS-161 and 431-QSSS-434, hSLC34A2 numbering). Binding of the negatively charged Pi is stabilized by the positive α-helical dipoles at the N-termini of HP1 and HP2 and partial charges from backbone amides and sidechain hydroxyls, especially contributed by the highly conserved Ser residues of the QSSS repeat motifs. This is reminiscent of how substrate selectivity works in the CLC family of chloride channels and transporters^30^. Specifically, Pi is coordinated by electrostatic interactions with the backbone amides of Ser160, Ser161, Ser433, and Ser434, and hydrogen bonding with the sidechains of Ser159, Ser161, Ser432, Ser434, Thr192 (TM2), and Thr465 (TM5) (Fig. 2C). T192K is known to cause the lung disorder PAM due to a lack of function of SLC34A2 (Fig. S13)^31^. Although the Gln residues from the QSSS motifs do not directly interact with the substrate, Gln158 forms a hydrogen-bond with Thr195 (part of the highly conserved TS-TNT motif), and Gln431 forms one with Thr468 (from the highly conserved SN--TTTT motif), contributing to a stabilization of the substrate binding site architecture. Notably, the mode of Pi substrate binding differs markedly between SLC34 importers and the human Pi exporter XPR1, which relies on positively-charged residues, primarily arginines, to mediate phosphate recognition^32^.

**Figure 2.**
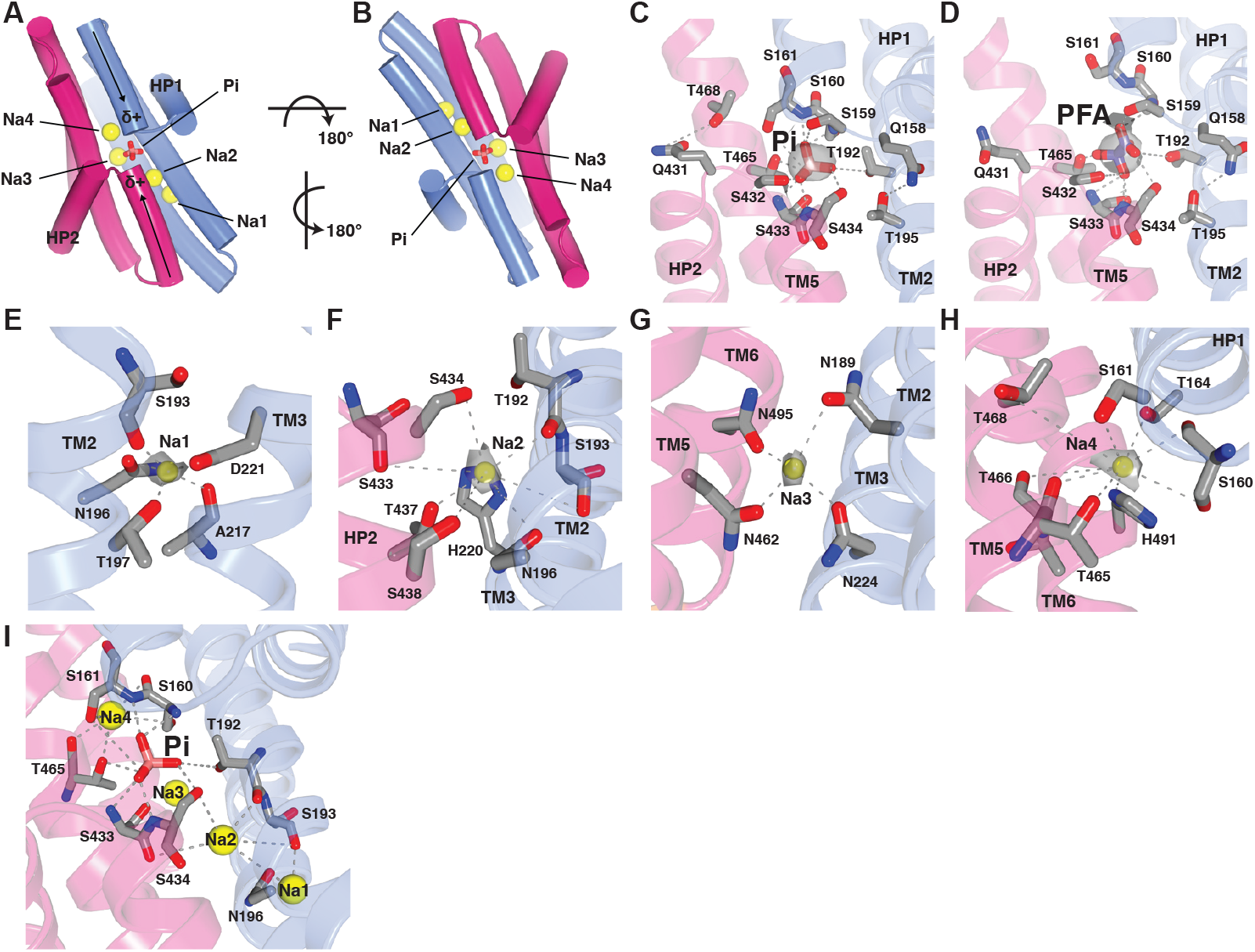
SLC34A2 substrate and ion binding. (A) Pi substrate (pink sticks) and Na^+^ ion (yellow spheres) binding to the transport domain of SLC34A2. (B) Symmetry of the Pi, Na2, Na3, and Na4 sites. (C) Interactions with the Pi substrate and the coordinating residues. The Pi is positioned by the positive helical dipole moments of HP1 and HP2. QSSS motif 1 consists of Gln158, Ser159, Ser160, and Ser161 and QSSS motif 2 consists of Gln431, Ser432, Ser433, and Ser434. Residue number corresponds to the human sequence. Pi density is depicted as a gray surface (1.5σ contour). Dashed lines indicate electrostatic and H-bond interactions. Nitrogen, blue; oxygen, red. (D) Interactions with the inhibitory substrate mimetic, PFA (purple sticks). PFA density is depicted as a gray surface (1.5σ contour). (E-H) Interactions with the Na^+^ ions (yellow spheres) and the coordinating residues for Na^+^-binding sites 1 (E), 2 (F), 3 (G), and 4 (H). Na^+^ ion densities are depicted as a gray surface (2.5σ contour (Na1 and Na3); 1.5σ contour (Na2 and Na4)). (I) Illustration of the interconnectedness between the Pi substrate and Na^+^ ions.

To determine the functional relevance of these observed interactions with the Pi substrate, we expressed human SLC34A2 (hSLC34A2) QSSS repeat motif mutants and T192A, T195A, T465A, and T468A in oocytes and measured Pi import activity using our radioactive [^32^P] transport assay. The expression of all mutants was detected using Western blot (Fig. S3). Serine residues were mutated to cysteine, expect for Ser159 and Ser432, which were mutated to alanine, as the S159C and S432C mutants were not expressed. All mutants have impaired transport, and Q158A, S161C, T192A, S434C, and T465A are nonfunctional (Fig. 3A). Reduced function of Q158A, T195A, Q431A, and T468A highlights the importance of the site’s unique architecture, particularly in the region where the helical hairpins intersect. Several analogous mutations in SLC34A1 have been shown to disrupt transporter function^33,34^, consistent with our findings and the notion that SLC34 family members have a shared substrate-binding site (Fig. S1).

**Figure 3.**
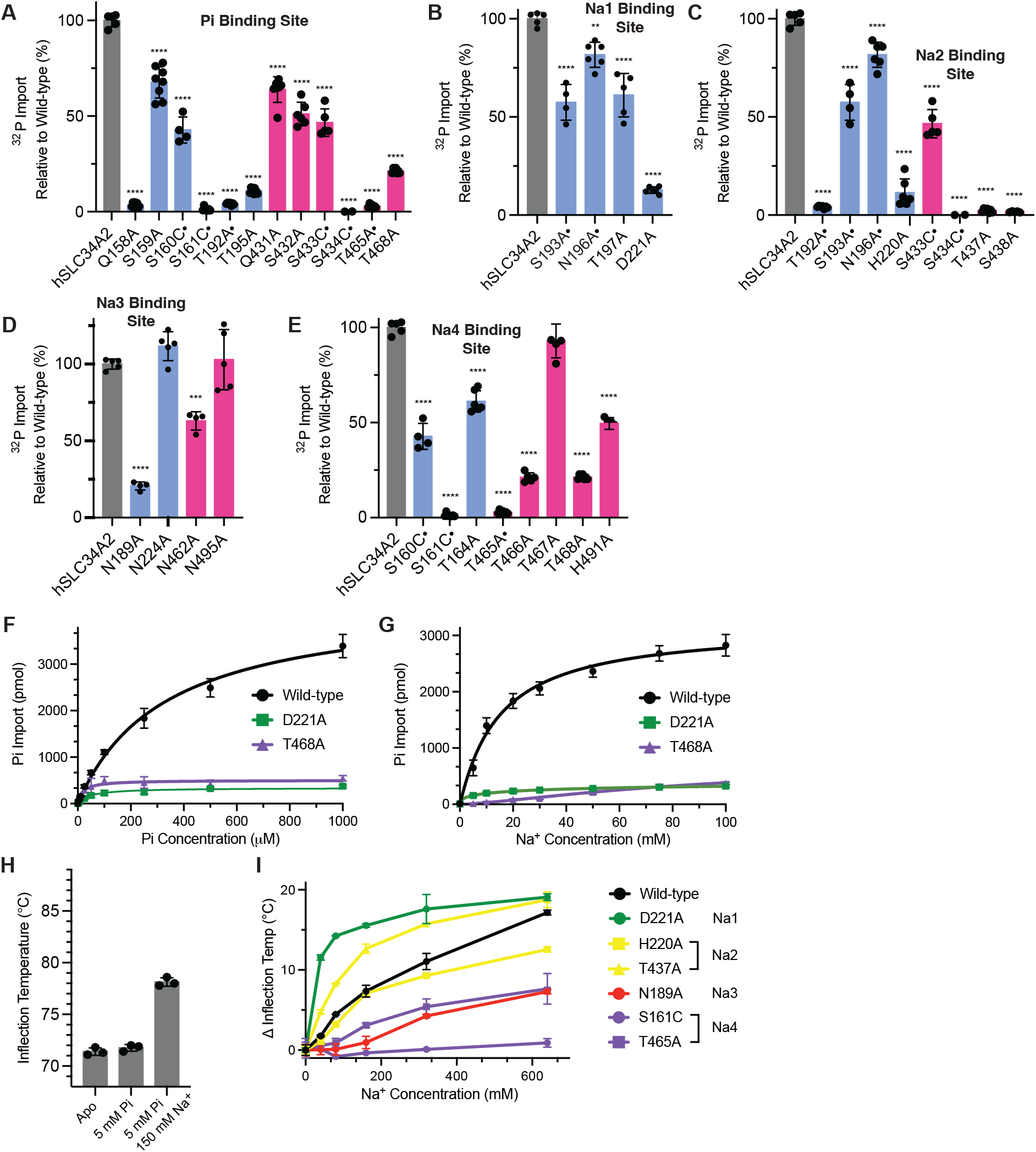
Functional analysis of the importance of and coupling between substrate and ion binding to SLC34A2. (A) Structure-function analysis demonstrating the importance of the QSSS motifs and neighboring Thr residues involved in Pi substrate binding. Experiments conducted in oocytes expressing hSLC34A2. Data are represented as the normalized mean ± SD (n = 4-9 samples). Significance was determined using one-way ANOVA (**** *P* < 0.0001). The dots denote residues that contribute to more than one substrate or ion binding site. (B-E) Structure-function analysis of the residues integral to Na^+^-binding sites 1 (B), 2 (C), 3 (D), and 4 (E). Data are represented as the normalized mean ± SD (n = 4-7 samples). Significance was determined using one-way ANOVA (** *P* < 0.0021, *** *P* < 0.0002, **** *P* < 0.0001). (F-G) Kinetic parameters of Na^+^-driven Pi transport by wild-type and mutant hSLC34A2 when Pi (F) and Na^+^ (G) concentrations are titrated. (H) Thermostability assays conducted with purified hSLC34A2 demonstrating that Pi requires Na^+^ binding to induce a change in the conformational landscape of the protein. (I) Evaluation of the impact of mutations that impair Na^+^ binding at each of the four sites on the conformational landscape of hSLC34A2 using nanoDSF.

### Sodium-binding sites

The SLC34A2-mediated uptake of Pi is driven by the Na^+^ gradient^2^. During the transport cycle, it has been proposed that two Na^+^ ions bind sequentially and cooperatively to facilitate access of Pi to its binding site^35,36^. After Pi binds, a third Na^+^ ion is thought to bind and reorient the fully loaded carrier, triggering transition to the occluded and inward-open states, thereby enabling substrate and ion translocation and their subsequent release into the cytosol. To test this model, we visualized Na^+^-binding sites by determining structures in the presence of 150 mM Na^+^ alone, as well as in the presence of Na^+^ and substrate. Unambiguous ion density was not observed in our Na^+^ alone map, most certainly due to its moderate 3.7 Å resolution. However, in our SLC34A2 maps in the presence of Na^+^ and Pi, we observed four discreate densities that are consistent with ion occupancy (Fig. 2A,B,E-H,I and Figs. S11C and S14). Assignment of these densities as Na^+^ was based on an integrated approach combining comparative analysis of multiple cryo-EM maps (-/+ Na^+^), sequence conservation across family members, intrinsic protein symmetry, evaluation of the local chemical environment, binding site prediction tools (Metric Ion Classification and AlphFold3^37,38^), the CheckMyMetal validation tool^39^, nanoDSF binding measurements, complementary functional assays, and agreement with established biological roles. We observed two Na^+^ ions (Na2 and Na4) that flank the Pi density and a third Na^+^ also in close proximity (Na3), as well as a fourth along the transport pathway but further towards the cytosolic side (Na1). Strikingly, the highly symmetric Na2, Pi, Na4, and Na3 binding sites are located at the interface between the two inverted repeats (Fig. 2A,B).

The first Na^+^ binding site sits away from the Pi substrate buried in a pocket formed by TM2 and TM3 and comprised of Ser193, Asn196, Thr197, Ala217, and Asp221 (Fig. 2E). SLC34A1 mutants with substitutions in the positions analogous to Ser193, Asn196, and Thr197 have both decreased transport activity and apparent substrate affinity^33,34,40^. The second Na^+^ binding site neighbors the Pi binding site and is comprised of both sidechain and backbone interactions with residues of HP2 (QSSS--TS motif – Ser433, Ser433, Thr437, and Ser438) and the TM2 and TM3 helices of inverted repeat 1 (TS-TNT motif – Thr192, Ser193, and Asn196) and His220 (Fig. 2F). Na^+^ binding site 3 is formed by a highly symmetric coordination through Asn189 and Asn224 from inverted repeat 1 and Asn462 and Asn495 from inverted repeat 2 (Fig. 2G). The fourth Na^+^ binding site once again neighbors the Pi binding site and is comprised of both sidechain and backbone interactions with residues from HP1 (QSSSTSTS motif – Ser160, S161, and Thr164) and from the TM5 and TM6 helices of inverted repeat 2 (TTTT motif – Thr465, Thr466, and Thr468) and His491 (Fig. 2H). The Pi substrate and Na^+^ ions communicate through an intricate network of interactions involving shared amino acid residues (Fig. 2I). For instance, the sidechain hydroxyl groups of Thr192 and Thr465 coordinate Pi and the backbone carbonyls of these residues coordinate Na2 and Na4, respectively. Notably, Na3 does not participate in this interaction network.

To explore potential functional roles for our four proposed Na^+^ binding sites, we analyzed Pi import for hSLC34A2 proteins bearing single alanine substitutions at the identified coordinating residues. Indeed, many of these mutants, including those targeting each of the four Na^+^ sites, show diminished transport function, validating their participation in SLC34A2 transport activity (Fig. 3B-E and Fig. S3). Thermal shift assays have proven to be a powerful approach for detecting substrate and ion binding to Na^+^-driven transporters^41^ and were therefore also used for purified hSLC34A2 to further probe the identified Na^+^-binding sites with high throughput. Upon recording thermal unfolding profiles, we observed that the addition of either 640 mM Na^+^ alone or 5 mM Pi and 150 mM Na^+^ increased the resistance of SLC34A2 to heat denaturation, with an average inflection temperature (ΔTi) increase of 6.7 °C and 4.8 °C, respectively, indicating molecular interactions between protein and substrates (Fig. 3H and S15A,B). When conducting Na^+^ titrations in the presence of 5 mM Pi, we observed protein stabilization at lower Na^+^ concentrations and a higher maximal ΔTi increase of 16.8 °C, suggesting further changes in protein structure when Pi is present (Fig. S15A-C). To further facilitate the confident assignment of these densities as Na^+^ ions and determine the contribution to thermostability of Na^+^ binding to individual sites, we analyzed thermostability shift assay responses for SLC34A2 transporters bearing substitutions at these sites (Fig. 3I and Fig. S12C). Mutants targeting sites Na1 (D221A) and Na2 (H220A and T437A) have more prominent or equivalent shifts in ΔTi. Na3 (N189A) and Na4 (S161C and T465A) mutants are characterized by substantially smaller ΔTi changes. These results suggest that the inability of Na^+^ to bind at specific sites stabilizes distinct intermediate states and emphasizes the interconnectedness of these sites. To further support our findings, we next calculated Na^+^ affinities using transport assays for hSLC34A2 transporters bearing alanine substitutions at two of the identified Na^+^-binding sites (Na1 binding mutant D221A and the Na4 binding mutant T468A) (Fig. 3G). The apparent Na^+^ affinity constant for wild-type transporter is 15 mM, which is similar to what has been measured previously^26-28^. Consistent with roles in binding Na^+^ ions, both mutants show a trend toward lower binding affinities. Together, the binding and transport assays underscore important roles for all four Na^+^ sites in SLC34A2 function.

Given that SLC34A2 transports its substrates with a 3:1 Na^+^ to substrate stoichiometry^42^, it had been assumed that SLC34A2 binds three Na^+^. Intriguingly, our work supports an alternative model wherein the binding of an additional, non-transported Na^+^ ion is also required. The fourth Na^+^ may remain stably bound or be released back into the extracellular space. Precedent for Na^+^ binding events that are not strictly coupled to translocation comes from studies of SLC34A3, which support a model in which three Na^+^ ions bind, while only two are transported alongside Pi^43^.

### The roles of sodium and coupling mechanisms

Secondary active transporters use distinct mechanisms to couple ion and solute fluxes, enabling them to meet specific physiological needs and environmental conditions. Comparison of our SLC34A2 structures in its apo state, bound to Na^+^ alone, and bound to both Pi and Na^+^ now allow us to define the distinct roles of Na^+^ ions in the transport cycle and how Pi and Na^+^ transport is coupled. Our structures predict that Na^+^ binds first to sites 1, 2, and 3, which helps create the Pi binding pocket. Pi then binds followed by a Na^+^ in site 4. Importantly, binding of Na4 facilitates the transition of the fully loaded SLC34A2 from the outward-open to the occluded state initiating the transport cycle.

In the apo structure, we observe helical unwinding midway through TM2, centered around Gly191 and Thr192 (Fig. 4A and Fig. S16A,D). This unwinding is stabilized by interactions between Asp221 and the backbone nitrogens of these two residues. Glycines often plays a disruptive role in α-helices, as the absence of a sidechain allows for greater conformational freedom leading to kinds, bends, and unwinding. This conformation is further stabilized by hydrophobic interactions between Ile194 and Phe537/Phe538, which reside within a π-helix on TM7 of the accessory domain. This π-helix may help position these sidechains to mediate critical interactions between the transport and accessory domains. Upon Na^+^ binding, Asp221 undergoes a substantial reorientation to permit coordination with Na1; concurrently, TM2 adopts a canonical α-helix conformation (Fig. 4B and Fig. S16B,C,E). These dramatic conformational changes in TM2 enable Na2 and Na3 binding, which requires repositioning of Thr192 and Asn189, respectively, residues originally located on opposite sides of the unwound TM2 region. Thr192 also directly interacts with the Pi substrate. These conformational changes were observed in both the Na^+^ alone and Pi- and Na^+^-bound structures (Fig. S16). Notably, the π-helix in TM7 is stabilized by Trp225 from TM3. Trp225 is wedged between Phe536 (a Cys residue in zfSLC34A2) and Phe537, and lies adjacent to Asn224, which participates in Na3 binding. This arrangement suggests that the accessory domain plays a key role in stabilizing distinct states throughout the transport cycle. The density for Na3 is the strongest. Given that Na3 is positioned away from the translocation pathway of Pi and the other Na^+^ ions, it is most likely to play a structural role in stabilizing SLC34A2 rather than being transported itself. Overall, Na1, Na2, and Na3 binding supports Pi binding to a pocket rich in partial negative charge and alters the local network of hydrogen bonds and salt bridges to yield a high-affinity Pi binding site configuration.

**Figure 4.**
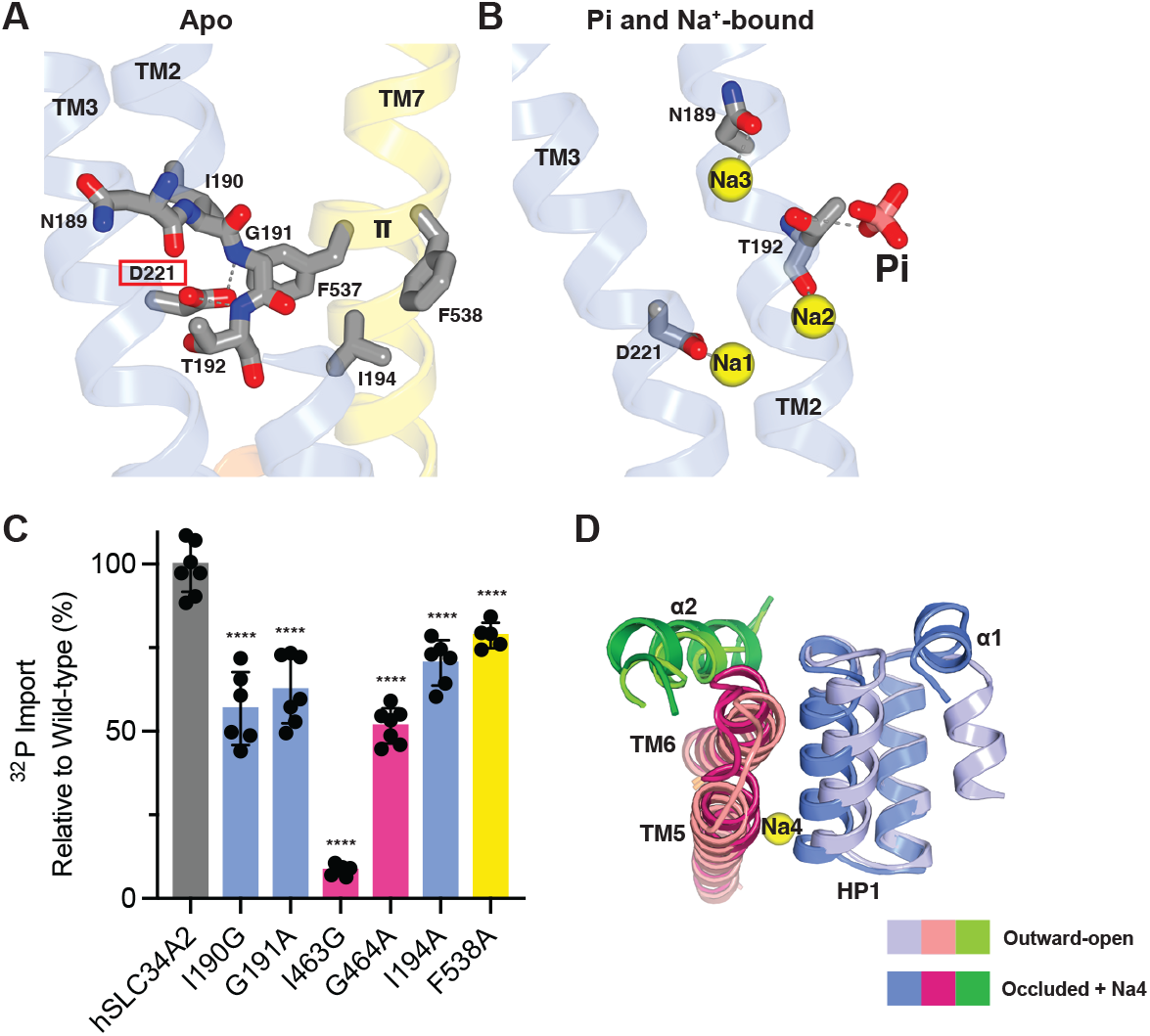
Role of Na^+^ ions in SLC34A2 Pi transport. (A) In the absence of Na1, TM2 contains a short-unwound stretch centered on Gly191 and stabilized by Asp221 and interactions with the accessory domain. (B) Upon Na1 binding, TM2 adopts a continuous α-helical conformation that facilitates Na2 and Na3 binding. Together, Na1, Na2, and Na3 form a favorable binding pocket for the Pi substrate. (C) Structure-function analysis testing the role of TM2 and TM5 unwinding for SLC34A2 transport. Data are represented as the normalized mean ± SD (n = 5-8 samples). Significance was determined using one-way ANOVA (**** *P* < 0.0001). (D) Upon Na4 binding, HP1 and TM5 are repositioned thereby closing the substrate entry pathway. This allows for a transition from the outward-open to the occluded state.

To test whether TM2 helical unwinding/restoration of the helical structure is critical to the formation of the Pi binding site, we mutated Gly191 to alanine. Alanine residues are helix stabilizers. We also mutated Ile190 to glycine to perhaps favor the unwound conformation. These mutants reduced transport activity by ∼50% (Fig. 4C), providing evidence for the importance of this structural feature. The inverted repeat topology of SLC34A2 prompted us to consider whether TM5 may also undergo unwinding at Gly464 and Thr465 during the transport cycle. Because Thr465 contributes to both the Pi and Na4 binding sites, unwinding of TM5 could potentially permit their release. In all our SLC34A2 structures, the entirety of TM5 is α-helical, however the synonymous mutations, I463G and G464A, also significantly reduce SLC34A2 function (Fig. 4C). The importance of these TM2 and TM5 regions is further emphasized by the nearby location of several PAM disease mutations, including G187E, T192K, and I198T in TM2 and G464R and T468del in TM5 (Fig. S13)^44^.

The final substrate to bind is Na4. Its coordination with key residues in HP1 and TM5 draws these two helices closer together, effectively repositioning them (Fig. 4D).

This movement is physically linked to conformational changes in the ECD, specifically altering the positions of the α1 and α2 helices. These shifts facilitate the transition of the transporter from an outward-open to an occluded state (further discussed below), highlighting the role of Na4 binding as a critical trigger for the structural rearrangements that drive the transport cycle.

To further assess how Na^+^ binding influences the alternating access cycle, we focused on two functionally distinct sites. Na1, which promotes formation of the substrate binding pocket, and Na4, which drives the transition to the occluded state. The site-specific mutants D221A (Na1) and T468A (Na4) exhibit markedly decreased apparent K_m_s for Pi (from 330 µM (95% confidence interval (CI): 290-380) to 47 µM (95% CI: 35- and 19 µM (95% CI: 14-27), respectively) (Fig. 3F). This suggests that disruption of Na^+^ coordination at either site stabilizes Pi high-affinity, non-productive states. D221A may permit substrate binding via TM2 rearrangement that bypasses Na1, whereas T468A may be impaired in driving the transition from the outward-open to the occluded state.

Although Pi and Na^+^ are not directly coordinated, Thr192, Ser433, and Ser434 also contribute to Na2 binding and Thr465, Ser160, and Ser161 also contribute to Na4 binding (Fig. 2I), suggesting functional coupling between substrate and these Na^+^ ions. Communication between Na^+^ sites 1 and 2 is facilitated by Ser193 and Asn196 on TM2, which interacts with both bound ions.

### Structural basis of alternating access

Our ensemble of structures defines two distinct conformational states of SLC34A2: (i) an outward-open state, which is most prevalent in the presence of Na^+^ only, when trapped by the antagonist PFA, or with a mutation in the first QSSS motif (S161C, S135C zebrafish numbering) that disrupts Na4 binding (Fig. 5A); and (ii) an occluded state, which is most prevalent in the absence of Pi and Na^+^ or in the presence of both (Fig. 5C). In the outward-open state, a cavity, created by a repositioning of the ECD, opens to the extracellular space and extends to the membrane center, such that Pi and Na^+^ have unhindered access to their binding sites (Fig. 5B). The electrostatic potential of the transporter surface creates a negatively charged extracellular region that attracts Na^+^ (Fig. S17).

**Figure 5.**
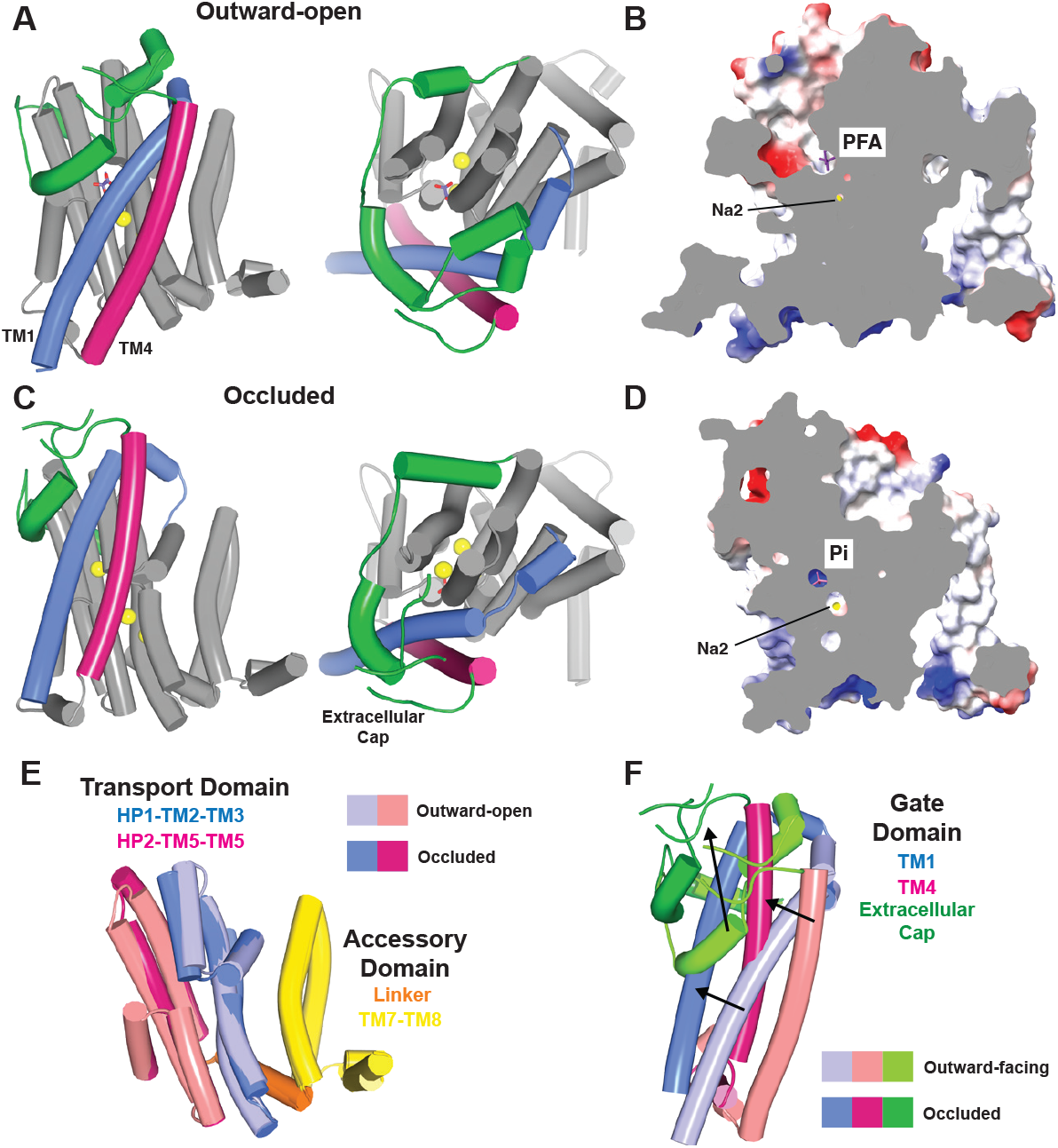
Conformational changes associated with transition from an outward-open to occluded state. ((A,C) Side and top views of SLC34A2 in the outward-open (A) and occluded (C) states. The gate domain is colored as in Fig. 1 with transmembrane helices shown as cylinders. Pi substrate (pink sticks), PFA (purple sticks), and Na^+^ ions (yellow spheres) are shown. (B,D) Cross-sectional surface representation of SLC34A2 in the outward-open (B) and occluded (D) conformations. (E) Superposition of the transport and accessory domains in the outward-open and occluded conformations. There is minimal movement, which is restricted to HP1 and TM5. (F) Superposition of the gate domain in the outward-open and occluded conformations. Arrows indicate the direction of the movement of the gate domain from the outward-open to occluded conformation.

Comparison of the outward-open and occluded conformations reveals the gating elements and structural rearrangements that drive alternating access of the substrate- and ion-binding sites in the SLC34A2 transport cycle. The transition from the outward-open to the occluded state involves two coordinated events, a subtle narrowing of the gap between HP1 and TM5 in the transport domain, facilitated by Na4 binding, and a pronounced upward repositioning of the entire gate domain. This latter movement encompasses the TM1 and TM4 helices, whose rearrangements are coupled to a rigid-body movement of the ECD, herein referred to as the extracellular cap, and in the intracellular α4 helix. Despite these large-scale motions typical of transporters, the SLC34A2 transport domain exhibits minimal conformational change (Figs 4D, 5E). In contrast, the peripheral gate domain adopts either a down position, characteristic of the outward-open state, or a neutral position, characteristic of the occluded state, correlated with substrate access. In the occluded state, the extracellular cap obstructs the entry pathway (Fig. 5F). The exceptionally dynamic nature of the gate domain likely accounts for its relatively poor resolution, as it samples a continuum of conformational states that are challenging to distinguish and separate during structural analysis.

The apical membranes of intestinal and renal epithelial cells, where SLC34 transporters reside, are enriched in cholesterol^45^. We observed a notable density for a cholesterol molecule at the interface between HP1 and TMs 1 and 4 in our occluded maps (Fig. S18). Cholesterol binds in a conformation-specific manner, as neither the lipid nor its binding site is present in the outward-open state. The observed shifts in TM1, TM4, and HP1, which define the gating mechanism, suggest that cholesterol binding may stabilize the occluded state and be critical for SLC34A2 transporter function.

### Mechansisms of inhibition and therapeutic antibody targeting

In CKD patients, the kidneys’ ability to excrete Pi is impaired, while Pi absorption in the intestine is increased, leading to elevated Pi levels throughout the body^46^. Inhibiting SLC34A2 to reduce dietary Pi uptake presents an attractive opportunity to modulate systemic Pi levels, a strategy that has garnered significant clinical interest ^4^. To explore mechanisms of SLC34A2 inhibition, we determined a structure of SLC34A2 in the presence of PFA and Na^+^. PFA, also known as foscarnet, is a pyrophosphate analogue that competitively blocks Pi uptake by SLC34 transporters^27,47,48^. The PFA antagonist resides in the Pi-binding site in a manner analogous to the Pi substrate, providing further support that this is indeed where substrate binds (Fig. 2D and Fig. S11B). This structure reveals how the PFA inhibitor arrests the SLC34A2 transport cycle. In the presence of PFA and Na^+^, we observed that only three of the four Na^+^ sites (Na1, Na2, and Na3) are occupied (Fig. S14E,F). This indicates that by blocking Na4 binding, PFA traps the transporter in its outward-open state and arrests the transport cycle. It also provides additional support for our mechanism that Na4 binding drives alternating access.

A humanized version of the SLC34A2-targeting MX35 antibody is currently in Phase II and Phase III clinical trials as an ADC to treat ovarian cancer. Our structures indicated that the large TM3-TM4 extracellular loop proposed to engage MX35 is highly flexible^49^. Although this loop cannot be fully modelled, we visualized its stabilization by a disulfide linkage between residues Cys277 and Cys321. Through determination of a low-resolution complex SLC34A2-MX35 structure, we have verified that therapeutic antibodies recognize an epitope within this loop^49^, and would decorate the surface of cancer cells (Fig. S2B).

## DISCUSSION

The movement of Pi underpins virtually every aspect of human physiology, from biomolecule biogenesis to cellular energy metabolism and signaling. The SLC34 family of Na^+^-dependent Pi importers mediates Pi uptake in key tissues, including the kidney and intestine, orchestrating systemic phosphate balance essential for health.

Members of the SLC34 family have long been proposed to operate via an elevator-type alternating access transport mechanism, in which the entire transport domain moves through the membrane as a rigid body to translocate substrate^50^. However, our data supports that the structural architecture and mechanisms of SLC34A2 alternating access fundamentally differ from classical elevator transporters (Fig. S16). First, SLC34A2 is a monomer, and most elevator transporters are multimers relying on a scaffold domain to facilitate the large movements of the transport domain. Instead, SLC34A2 may utilize its amphipathic linker helix and accessory domain to provide intrinsic structural support, thereby stabilizing the transporter within the membrane. Second, the transport domain and substrate- and ion-binding sites in SLC34A2 remain primarily stationary during conformational transitions. Instead, SLC34A2 gates via a highly mobile peripheral domain that traverses the membrane to mediate alternating access. The relatively moderate to slow transport rate of SLC34s is consistent with the substantial conformational dynamics required for their alternating access mechanism^28,51^. Third, an extracellular cap domain prevents substrate access in the occluded state. The SLC34 mechanism does not conform to canonical transporter classifications of rocker-switch, rocking-bundle, or elevator^52^, but instead exemplifies an atypical variation of the elevator-type mechanism. Similar gating behavior has been observed in the calcium/cation antiporter family, although these transporters are trimeric^53-55^. Our results support an emerging concept in which the mobile elevator domain of a transporter can either directly translocate substrates or modulate their movement indirectly.

The intrinsic inverted pseudosymmetry of SLC34A2 enables us to infer the likely architecture of its inward-open state. This is because the helical hairpins and transmembrane helices that coordinate and gate the substrate and ions from the extracellular side are structurally related, a feature commonly observed in transporters and exploited to mediate alternating access ^56^. Therefore, we postulate that transition to the inward-open state involves an upward displacement of the peripheral gate domain. Furthermore, local unwinding of TM5 may facilitate Pi release by disrupting the coordination network within the binding pocket. In contrast to the extracellular cap, which restricts access from the outside, no corresponding intracellular cap is apparent, suggesting that cytosolic release is less sterically constrained, although the α4 helix may function as an intracellular gate. Ultimately, visualization of all major functional states will be required to unequivocally reveal the complete transport cycle.

SLC34A2 transport is electrogenic, resulting in one net positive charge transported per cycle^23^. The SLC34A3 Pi transporter couples one less Na^+^ ion to Pi transport, in comparison to SLC34A2 and A1^35^. Our structures provide a mechanistic rationale as to why. While the Na2, Na3, and Na4 binding sites are conserved amongst all SLC34 family members, there are differences in Na^+^ binding site 1 with a key change being the substitution of Asp221 for a glycine residue and Asn196 for a serine residue (Fig. S1). This is consistent with previous reports that SLC34A3 transport becomes electrogenic if three amino acids in this region that differ in both charge and polarity are substituted (S190A, S192A, G196D corresponding to Ala215, Ala217, and Asp221 in SLC34A2)^42^. Our work hints at an adaptation in SLC34A3 that tunes the transporters energy landscape making Na1 binding dispensable^57^ and thereby altering the transporter stoichiometry. Further investigation is warranted to link the molecular properties of SLC34 family members to their critical and targeted roles in health and disease.

### Data, Materials, and Software Availability

Cryo-EM density maps have been deposited to the Electron Microscopy Data Bank (EMDB) under the accession numbers EMD-73829 (apo zfSLC34A2), EMD-73830 (Na^+^-bound zfSLC34A2), EMD-73831 (Pi- and Na^+^-bound zfSLC34A2), EMD-73832 (PFA- and Na^+^-bound zfSLC34A2), EMD-738233 (apo zfSLC34A2 S135C), EMD-73834 (Pi- and Na^+^-bound zfSLC34A2 S135C). Atomic coordinates have been deposited in the Protein Data Bank (PDB) under IDs 9Z60 (apo zfSLC34A2), 9Z61 (Na^+^-bound zfSLC34A2), 9Z62 (Pi- and Na^+^-bound zfSLC34A2), 9Z63 (PFA- and Na^+^-bound zfSLC34A2), 9Z64 (apo zfSLC34A2 S135C), 9Z66 (Pi- and Na^+^-bound zfSLC34A2 S135C). All other data are included in the manuscript and/or SI Appendix.

## Acknowledgments

We thank K. Enquist for her contribution to establishing the radioactive transport assay; F. Weis-Garcia and the staff of the Antibody and BioResource (ABR) Core at Memorial Sloan Kettering Cancer Center (MSKCC) (RRID: SCR_017691) with whom the zfSLC34A2 monoclonal antibody was generated; M. J. de la Cruz and S. Sen of the Structural Biology core facility at MSKCC (RRID: SCR_027809) for help with cryo-EM data acquisition; and A. Dar, R. Hite, D. Julius and members of the Diver laboratory for discussions. NanoDSF experiments were performed in the Molecular Biophysics core facility at Weill Cornell Medicine under the leadership of R. Rusinova. M.M.D. is a Josie Robertson Investigator. This research is supported in part through the National Institutes of Health (NIH) / National Cancer Institute (NCI) Cancer Center Support Grant P30CA008748.

## Author contributions

Q.Z. and M.M.D. designed and executed experiments, including protein expression and purification, monoclonal antibody generation, cryo-EM data acquisition and image processing, atomic model building and refinement of SLC34A2 structures, as well as binding assays and radioactive transport assays. O.A. was instrumental in the initial cloning, ortholog screening, and optimization of the purification protocol and transport assay. M.M.D. wrote the manuscript, with input from all authors.

## MATERIALS AND METHODS

### Cloning, expression, and purification of SLC34A2

zfSLC34A2 was selected as a candidate for protein purification and structure determination using fluorescence-detection size-exclusion chromatography (FSEC) screening^58^. The full-length human and zebrafish SLC34A2 cDNA (synthesized by Twist Biosciences) was cloned into a modified pEG BacMam expression vector (addgene, 160686) containing an N-terminal Strep tag (WSHPNFEK) and mCerulean and followed by a Precision Protease cleavage site (LEVLFQ/GP) to facilitate its removal. To allow for MX35 Fab binding, we engineered a construct in which the MX35 epitope (SPSLCWTDGIQNWTM), which is not conserved between the human and zebrafish orthologs was inserted (zebrafish residues 295-300 were swapped for human residues 324-338), and this construct was used for structure determination.

SLC34A2 was expressed in Expi293F cells using transient transfection. Briefly, 100 µg of plasmid DNA was incubated with 300 µg of polyethylenimine (PEI 25,000) (Polysciences, 23966-1) for 30 min at room temperature, then added to 100 ml of cells at a density of 3.0x10^6^ cells/ml. Cells were supplemented with 10 mM sodium butyrate after 20 h, harvested 48 h post-transfection, and stored at -80°C.

To prepare sample for antibody generation, a cell pellet of zfSLC34A2 was re-suspended in buffer containing 25 mM Tris, pH 7.5, 200 mM NaCl, a cOmplete Protease Inhibitor Cocktail Tablet (Roche). 1% glycol-diosgenin (GDN, Anatrace, GDN101) was added to the cell lysate and the mixture was rotated at 4°C for 1 h to extract SLC34A2 from membranes. The sample was centrifuged at 50,000 x g for 50 min at 4°C and the supernatant was filtered through a 0.22-μm polystyrene membrane (Millipore Sigma). The sample was incubated with Strep-Tactin XT 4Flow high capacity resin (IBA Lifesciences, 2-5030-002) at 4°C for 1 h with rotation. At room temperature, beads were collected on a column, washed with buffer containing 25 mM Tris, pH 7.5, 200 mM NaCl, 0.002% GDN, and the protein as eluted with an identical buffer supplemented with 10 mM Biotin (IBA Lifesciences, 2-1016-002). The N-terminal Strep/mCerulean tag was removed by rotating the sample overnight at 4°C with PreScission Protease. SLC34A2 was further purified using a Superose 6 Increase size-exclusion column (SEC) (Cytiva) in 25 mM Tris, pH 7.5, 200 mM NaCl, 0.002% GDN.

To prepare sample for cryo-EM structure determination, the MX35 swapped zebrafish SLC34A2 proteins (both wild-type and S135C mutant) were expressed and purified in a manner identical. Before purification by SEC, 12H07-Fab was added at a 1:1.2 ratio (SLC34A2:12H07). The complex was then incubated on ice for 30 mins. Peak fractions were pooled and concentrated to 10-20 mg/ml using a 100-kDa concentrator (Amicon Ultra, Millipore Sigma). For the apo (K^+^) sample, 200 mM KCl replaced the NaCl in all purification buffers. For the Na^+^/Pi sample, 5 mM Pi was added in all purification buffers. For the Na^+^/PFA sample, 10 mM PFA was added in all purification buffers. For the MX35-bound sample, before purification by SEC, MX35-Fab was added at a 1:1.2 ratio (SLC34A2:MX35). The complex was then incubated on ice for 30 mins. Peak fractions were pooled and concentrated to 3.5 mg/ml.

To prepare sample for thermal stability analysis via nanoDSF, the hSLC34A2 protein and mutants were expressed and purified in a manner identical to apo zfSLC34A2 in the absence of ion and substrate.

### Antibody generation and Fab production

Monoclonal antibodies were raised in mice by the Antibody and BioResource (ABR) Core of the Memorial Sloan Kettering Cancer Center (MSKCC). zfSLC34A2 purified in GDN was used as the antigen for immunization. The antibody selection process included enzyme-linked immunosorbent assay (ELISA) and FSEC analysis to identify antibodies that preferentially bound to native SLC34A2 in comparison to SDS-denatured protein. This selection process yielded 12H07. 12H07-IgG2b was expressed using moue hybridoma cells, purified using Protein G Resin (GenScript, L00209), and cleaved using papain (Pierce Fab Preparation Kit, Thermo Fisher Scientific, 44985) to generate the Fab fragment. The Fab fragment was purified by SEC (Superdex 200 Increase column, Cytiva) in a buffer containing 25 mM Tris, pH 7.5, and 200 mM KCl.

### Cryo-EM sample preparation and data collection

Cryo-EM grids were prepared using a FEI Vitrobot Mark IV (Thermo Fisher Scientific) by applying 3 μl of purified XPR1 to a glow-discharged holy carbon gold QUANTIFOIL Au 1.2/1.3 grid (400 mesh, Electron Microscopy Sciences) and blotting for 1 s at 4 °C and 100% humidity, then plunge freezing in liquid ethane. Cryo-EM datasets were collected on FEI Titan Krios microscopes (Thermo Fisher Scientific) operated at 300 kV housed in the MSKCC Richard Rifkind Center for Cryo-EM. Images were recorded in an automated fashion on a K3 Summit Direct Electron Detector (Gatan) in super-resolution counting mode with a super-resolution pixel size of 0.413 Å (physical pixel size of 0.826 Å) at a dose rate of 15 e-/pixel/s using SerialEM or a Falcon 4i Direct Electron Detector (Thermo Fisher) with a Selectris X Energy Filter (Thermo Scientific) in counting mode with a physical pixel size of 0.725 Å at a dose rate of 11.6 e-/pixel/s using Smart EPU. The defocus range was − 0.7 to − 1.7 μm. Data collection statistics are shown in Supplementary Table S1.

### Electron microscopy data processing

Image processing was performed using cryoSPARC (v4.5.0 or v4.5.1)^59^. The videos were gain-corrected, Fourier-cropped by two (0.826 Å) (when using the K3 camera only), and aligned using whole-frame and local-motion-correction algorithms within cryoSPARC. Phenix Resolve CryoEM was to improve the maps through density modification and to provide FSC calculations and reliable resolution estimation^60^.

All the datasets were processed using a similar cryo-EM workflow, as summarized in Figs. S4-9. Blob-based auto-picking was implemented to select initial particles. Several rounds of two-dimensional (2D) classification were carried out, and the best 2D classes (those clearly bound to Fab) were manually selected to generate an initial three-dimensional (3D) model. Particles with easy-to-identify transmembrane helices were used for Topaz training and particle picking using neural networks. False-positive selections and contaminants were excluded through iterative rounds of heterogenous classification using the model generated from the ab initio algorithm as well as several decoy classes representing noise. 3D classification using a focus mask covering SLC34A2 and the Fv portion only of the 12H07 density was used to separate well-aligning particles. The structures were refined using non-uniform refinement and local refinement, followed by reference-based motion correction to generate the final reconstructions.

### Model building, refinement, and validation

ModelAngelo was used to automatically build initial atomic models into the cryo-EM density maps^61^. Subsequently, the models were manually rebuilt using Coot to improve the fit within the density ^62^. Atomic coordinates were refined against their corresponding density-modified maps using PHENIX real space refinement with geometric and Ramachandran restraints maintained throughout^63^. Model validation was performed using MolProbity^64^. Molecular graphics Figs. were prepared using UCSF Chimera^65^ or PyMOL. For electrostatic calculations, the APBS plugin in PyMOL was used^66^.

### Radioactive Pi transport assays

Radioactive ^32^P substrate influx experiments were carried out as follows. For transporter expression, *Xenopus laevis* oocytes (EcoCyte BioScience) were injected with 25 ng complementary RNA (cRNA) (synthesized using the mMESSAGE mMACHINE T7 Transcription Kit (Thermo Fisher Scientific, AM1344) encoding wild-type or mutant SLC34A2 transporters and incubated at 18°C in ND96 with calcium and antibiotics (5 mM HEPES, pH 7.4, 96 mM NaCl, 2 mM KCl, 1 mM MgCl_2_, 1 mM CaCl_2_, 50 µg/mL gentamicine, and 50 µg/mL tetracycline) for 3 days. For the transport assay, the oocytes were incubated in ND96 with calcium (5 mM HEPES, pH 7.4, 96 mM NaCl, 2 mM KCl, 1 mM MgCl_2_, 1 mM CaCl_2_) containing 1 mM cold Pi and ^32^P (specific activity 10 mCi mmol^−1^ Pi) for 20 mins at room temperature. Then, the cells were washed four times in cold ND0 with calcium and Pi (5 mM HEPES, pH 7.4, 100 mM choline chloride, 2 mM KCl, 1 mM MgCl_2_, 1 mM CaCl_2_, 2 mM Pi) and subsequently lysed using 600 µL of 2% SDS (Fisher Scientific, BP8200100). Samples were transferred to a scintillation vial with 5 mL of scintillation cocktail (Perkin Elmer, 6013327). Radioactivity was measured with a liquid scintillation analyzer (Perkin Elmer, Tri-Carb 2910-TR). Influx is graphed as pmol of Pi or for the mutants as a percentage relative to wild-type using Graphpad Prism 10. All transport assay data are reported as means ± standard deviation from 6-10 replicates. Significance was determined using one-way ANOVA (** *P* < 0.0021, *** *P* < 0.0002, **** *P* < 0.0001).

For experiments where the concentration of Pi was varied (0-1 mM), 96 mM NaCl was used. For the experiments where the concentration of NaCl was varied (0-100 mM), 1 mM Pi was used, and choline chloride was supplemented to ensure a constant cation concentration. For initial velocity curves, the data were fit to the following models using least squares nonlinear regression with GraphPad Prism 10 software. 1) For the initial velocity versus Pi concentration curves, a Michaelis-Menten model was used. 2) For the initial velocity versus Na^+^ concentration curves, an allosteric sigmoidal model was used.

To test whether 12H07 Fab binding alters SLC34A2 transporter activity, 50 ng of 12H07 Fab (∼1 mM) was injected into mCer-zfSLC34A2-MX35 expressing oocytes 4 h prior to beginning the transport assay.

For FSEC and Western blot detection to assess SLC34A2 construct expression, 3 injected oocytes were pooled. The oocytes were lysed in 150 µL of buffer containing 25 mM Tris-HCl (pH 7.5), 200 mM NaCl, cOmplete Protease Inhibitor Cocktail, 1% GDN by pipetting then rotating at 4°C for 1 h. The sample was centrifuged at 18,200 x g for 1 h at 4°C to pellet cell debris. The supernatant was then subject to FSEC analysis or separated by SDS-PAGE and transferred for Western blot analysis using an iBlot 2 NC stack (Thermo Scientific; 1B23001). Membranes were blocked in TBST + 10% milk, then incubated for 1h at room temperature with 0.5 µg/mL MX35 antibody (MSK ABR Core) and an anti-Δ-actin (Santa Cruz Biotechnology, sc-47778). An HRP-conjugated anti-mouse secondary antibody (1:5000 dilution, 1 h, room temperature) (Promega, W402B) was used for visual the bands. β-actin is the loading control. This measures total protein and cannot distinguish plasma membrane from intracellular transporter, leaving open the possibility that surface trafficking differences contribute to the functional effects.

### Thermostability analysis

Wild-type hSLC34A2 and its mutants were purified in a buffer containing 25 mM Tris-HCl (pH 7.5), 200 mM KCl, 0.002% GDN, then diluted to 3 µM in the same buffer. 10 µL of purified protein was combined with 10 µL reaction buffer containing 25 mM Tris-HCl, pH 7.5, 0.002% GDN, 10 mM Pi, 0, 80, 160, 320, 640, or 1280 mM NaCl and 1280, 1200, 1120, 960, 640, or 0 mM KCl to ensure maintain identical cation concentrations. The samples were run on a Tycho NT.6 system in Tycho capillaries (both by NanoTemper Technologies; capillaries TY-C001). The platform runs experiments which utilize Tycho technology, a modified version of nano Differential Scanning Fluorimetry (nanoDSF). The Tycho NT.6 runs a thermal ramp from 35°C to 95°C at a defined rate of 30°C/min and records intrinsic protein fluorescence from tryptophan and tyrosine residues at 330 nm and 350 nm throughout the run. The brightness ratio 350 nm / 330 nm is plotted against the temperature in the sample, resulting the unfolding profile. DSFworld was used to analyze the to calculate inflection temperatures (T_i_), a measure for the temperature(s) at which the protein unfolds^67^.

