## Supporting Information for "Structural and mechanistic insights into SLC34 phosphate import"

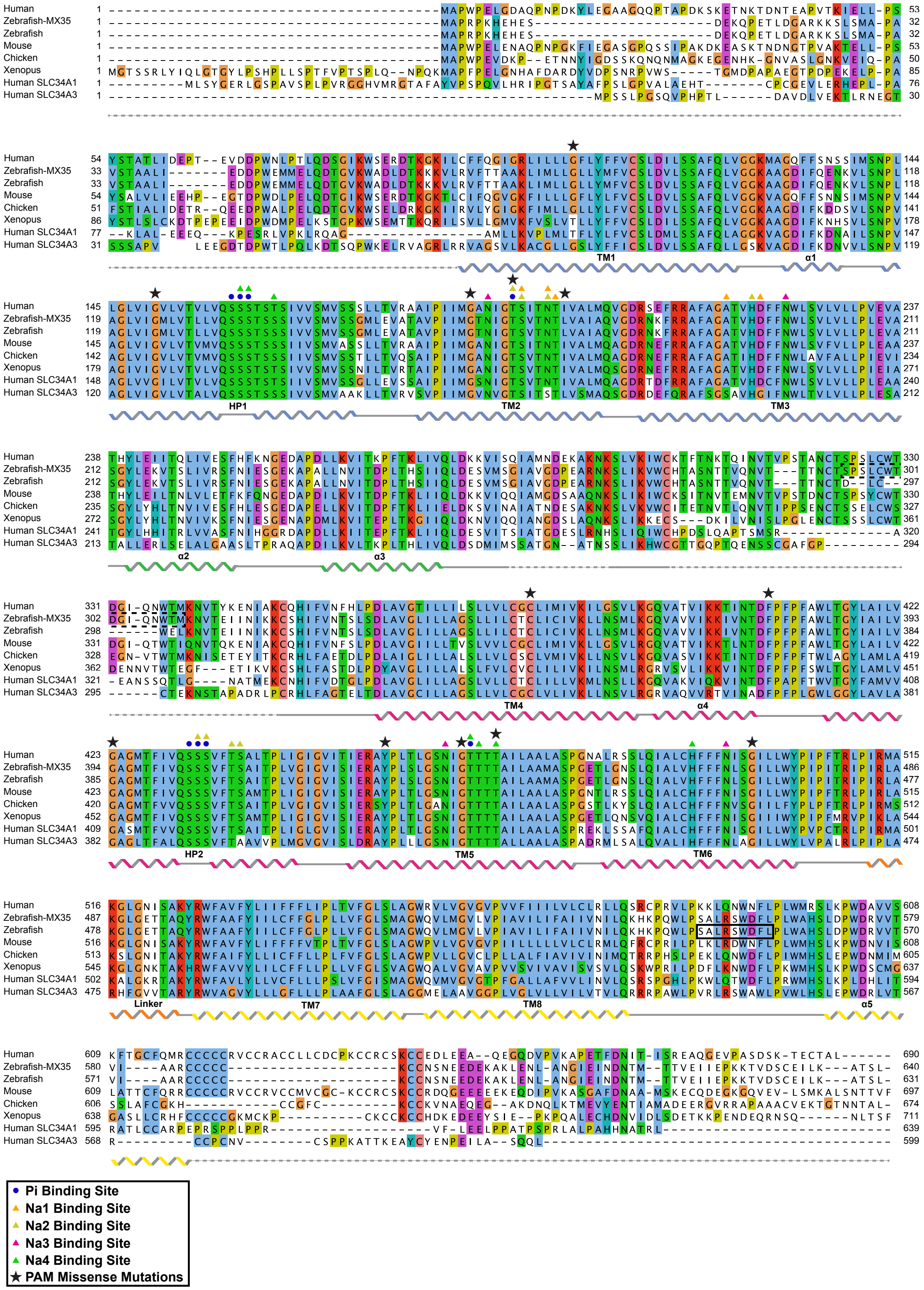


**Fig. S1.** Sequence alignment of SLC34 family members and SLC34A2 orthologs.

The protein sequences of SLC34A2 from human (Uniprot – O95436), zebrafish MX35, zebrafish (Uniprot - Q9PTQ8), mouse (Uniprot – Q9DBP0), chicken (Uniprot – A0A8V0YATA (first 10 residues deleted)), and frog (Uniprot – A0A8J0SQD6), SLC34A1 from human (Uniprot – Q06495), and SLC34A3 from human (Uniprot – Q8N130) were aligned using the Clustal Omega server. The secondary structure is indicated with ribbons representing $\alpha$-helices, solid lines representing loop regions, and dashed lines representing disordered regions. Assignments are based on the occluded (Pi- and $Na^+$-bound) structure. The alignment is colored according to the ClustalW convection. The solid black box indicates where the 12H07 Fab binds. The dashed black box indicates where the MX35 therapeutic antibody binds. SLC34A2 residues identified in this study as important for Pi and $Na^+$ binding are indicated. Black stars mark known PAM disease-causing missense mutations.

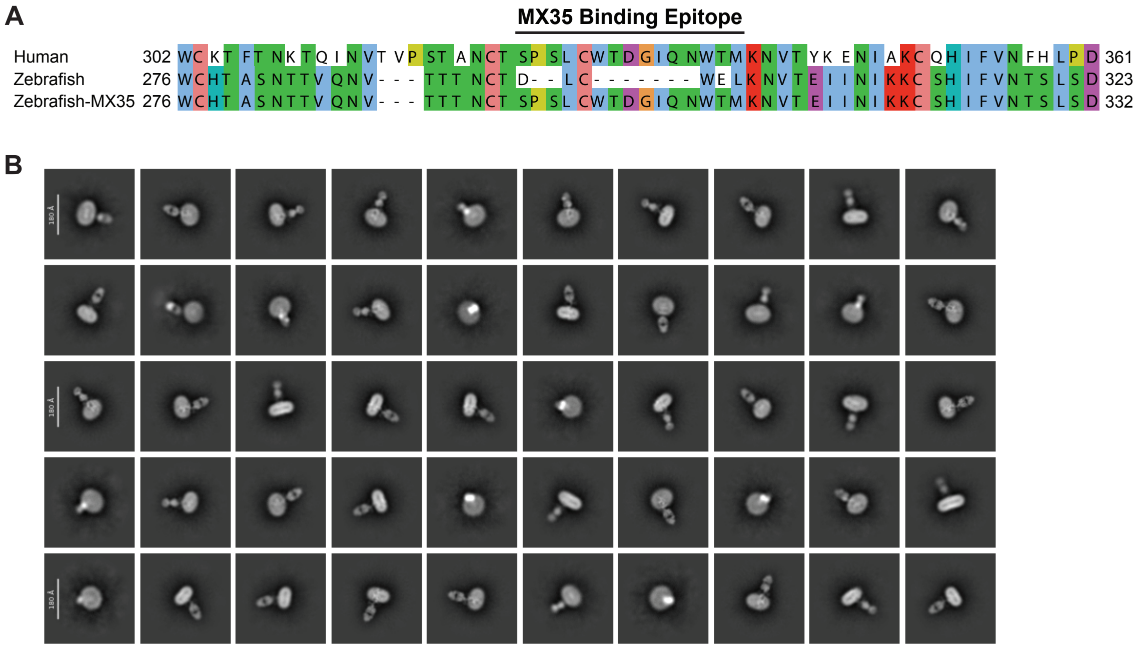


**Fig. S2.** Binding of the therapeutic MX35 antibody.

(A) Sequence alignment of the MX35 binding epitope within the flexible ECD. The binding site is not conserved with the zebrafish ortholog, therefore we engineered a construct, zfSLC34A2-MX35, with the epitope inserted. (B) Representative 2D class averages from cryo-EM analysis of the zfSLC34A2-MX35 + MX35 Fab complex. The gap between the detergent-embedded SLC34A2 and the MX35 Fab highlights the mobility of the ECD.

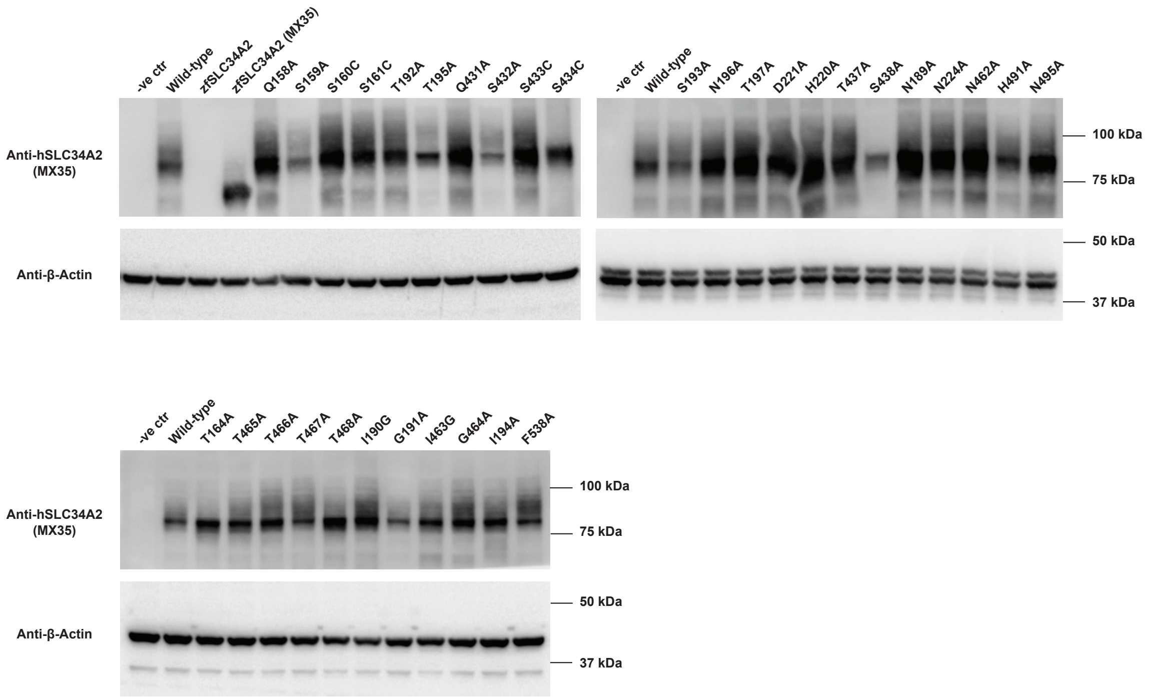


**Fig. S3.** Expression levels of SLC34A2 constructs.

Western blot analysis of *Xenopus* oocytes expressing wild-type or mutant SLC34A2. All mutants for which transport activity is measured are expressed. The negative control (-ve ctr) is uninjected oocytes. β-actin serves as a sample processing control.

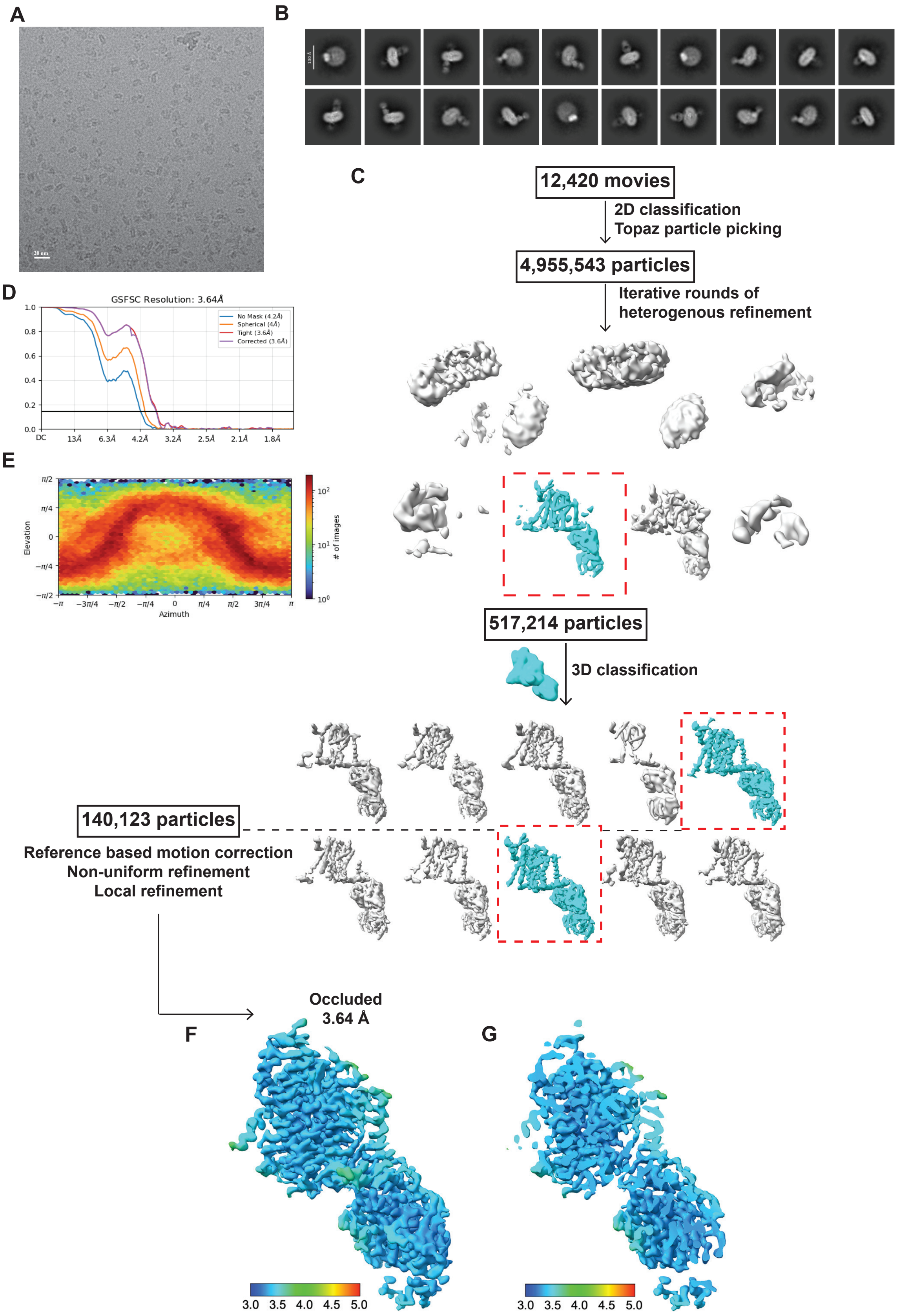
A
B
C
12,420 movies
2D classification
Topaz particle picking
4,955,543 particles
Iterative rounds of
heterogenous refinement
517,214 particles
3D classification
140,123 particles
Reference based motion correction
Non-uniform refinement
Local refinement
D
GSFSC Resolution: 3.64Å
No Mask (4.2Å)
Spherical (4Å)
Tight (3.6Å)
Corrected (3.6Å)
E
Elevation
Azimuth
### of images
F
Occluded
3.64 Å
G
3.0 3.5 4.0 4.5 5.0
3.0 3.5 4.0 4.5 5.0

**Fig. S4.** Cryo-EM data processing of apo zfSLC34A2.

(A) Representative micrograph. (B) Representative 2D class averages. (C) Flowchart for cryo-EM data processing. (D) Gold-standard Fourier shell correlation (FSC) curve (cutoff of 0.143) of the final density map. (E) Angular orientation distribution of all particles used in the final 3D reconstruction. (F) Final density map used for structural modelling. Surface color indicates estimated local resolution. (G) A cut-open view.

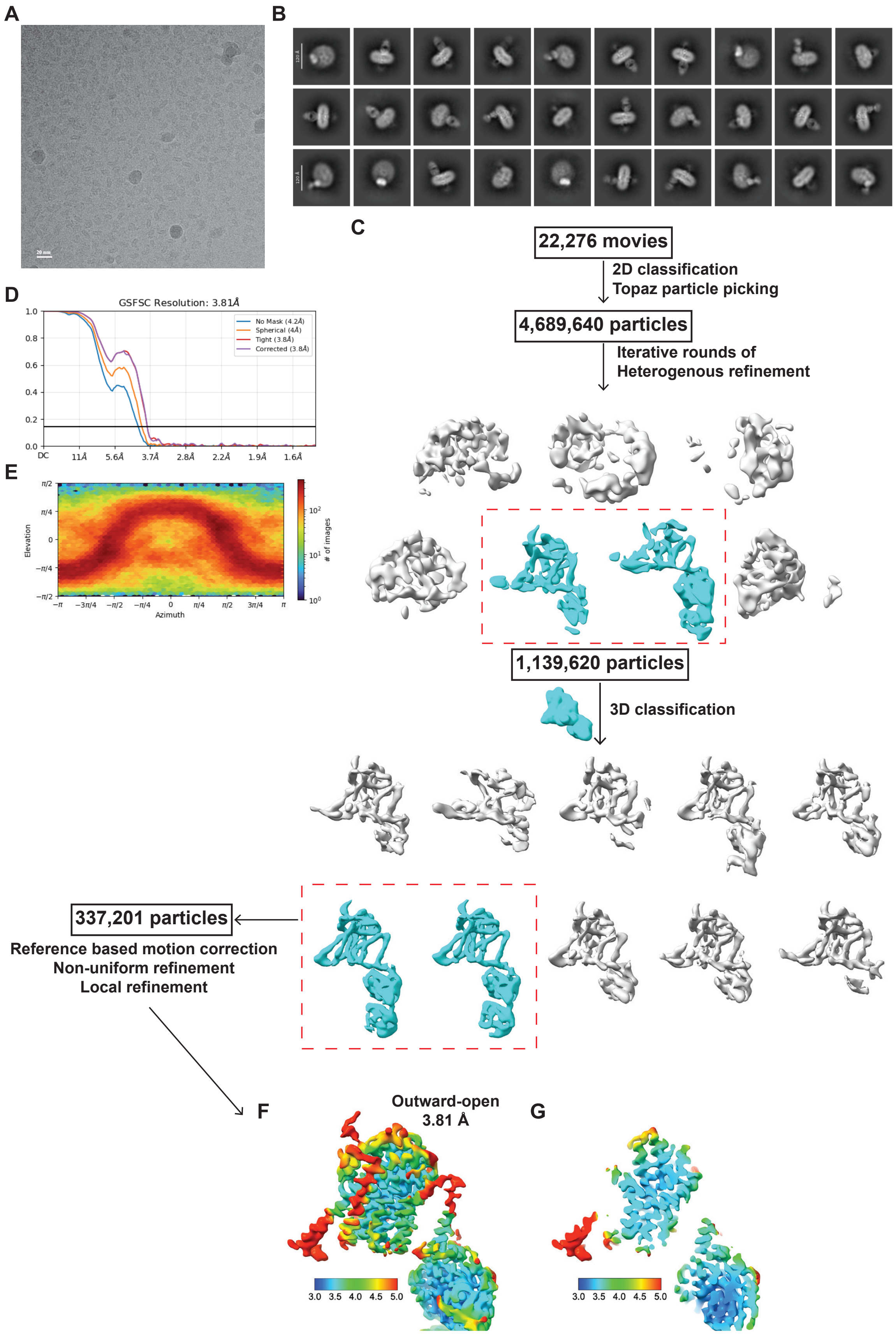
A
B
C
22,276 movies
2D classification
Topaz particle picking
4,689,640 particles
Iterative rounds of
Heterogenous refinement
1,139,620 particles
3D classification
337,201 particles
Reference based motion correction
Non-uniform refinement
Local refinement
D
GSFSC Resolution: 3.81Å
No Mask (4.2Å)
Spherical (4Å)
Tight (3.8Å)
Corrected (3.8Å)
DC
11Å
5.6Å
3.7Å
2.8Å
2.2Å
1.9Å
1.6Å
E
Elevation
Azimuth
### of images
F
Outward-open
3.81 Å
G
3.0 3.5 4.0 4.5 5.0

**Fig. S5.** Cryo-EM data processing of $Na^+$-bound zfSLC34A2.

(A) Representative micrograph. (B) Representative 2D class averages. (C) Flowchart for cryo-EM data processing. (D) Gold-standard FSC curve (cutoff of 0.143) of the final density map. (E) Angular orientation distribution of all particles used in the final 3D reconstruction. (F) Final density map used for structural modelling. Surface color indicates estimated local resolution. (G) A cut-open view.

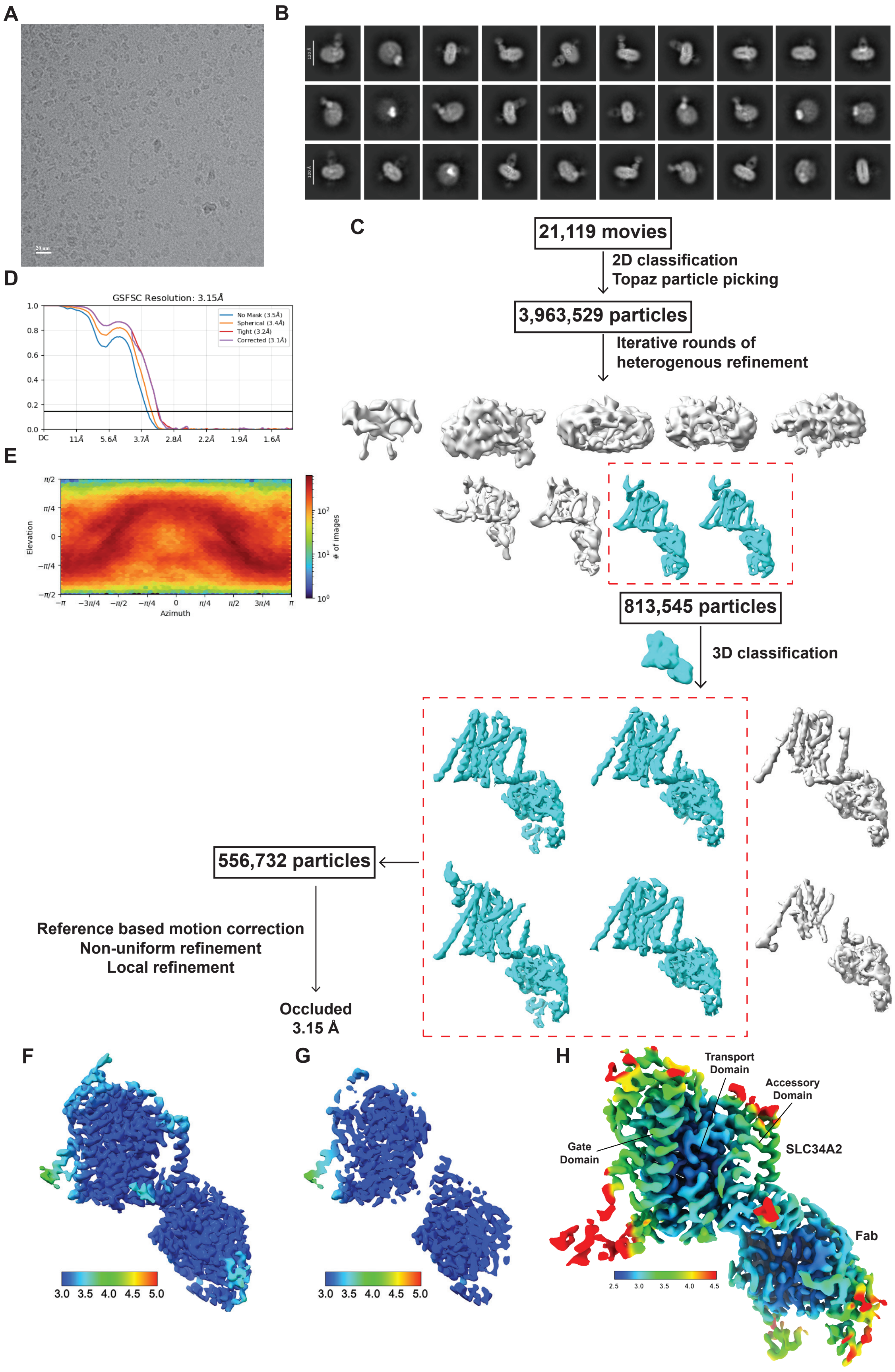

A
B
C
21,119 movies
2D classification
Topaz particle picking
3,963,529 particles
Iterative rounds of
heterogenous refinement
813,545 particles
3D classification
556,732 particles
Reference based motion correction
Non-uniform refinement
Local refinement
Occluded
3.15 Å
D
GSFSC Resolution: 3.15Å
No Mask (3.5Å)
Spherical (3.4Å)
Tight (3.2Å)
Corrected (3.1Å)
E
Elevation
Azimuth
### of images
F
G
H
Transport Domain
Accessory Domain
Gate Domain
SLC34A2
Fab

**Fig. S6.** Cryo-EM data processing of Pi- and $Na^+$-bound zfSLC34A2.

(A) Representative micrograph. (B) Representative 2D class averages. (C) Flowchart for cryo-EM data processing. (D) Gold-standard FSC curve (cutoff of 0.143) of the final density map. (E) Angular orientation distribution of all particles used in the final 3D reconstruction. (F) Final density map used for structural modelling. Surface color indicates estimated local resolution. (G) A cut-open view. (H) Cryo-EM map colored by local resolution, highlighting variation across domains. The highest resolution (~2.5 Å) is achieved in the transport domain where the substrate and ions bind.

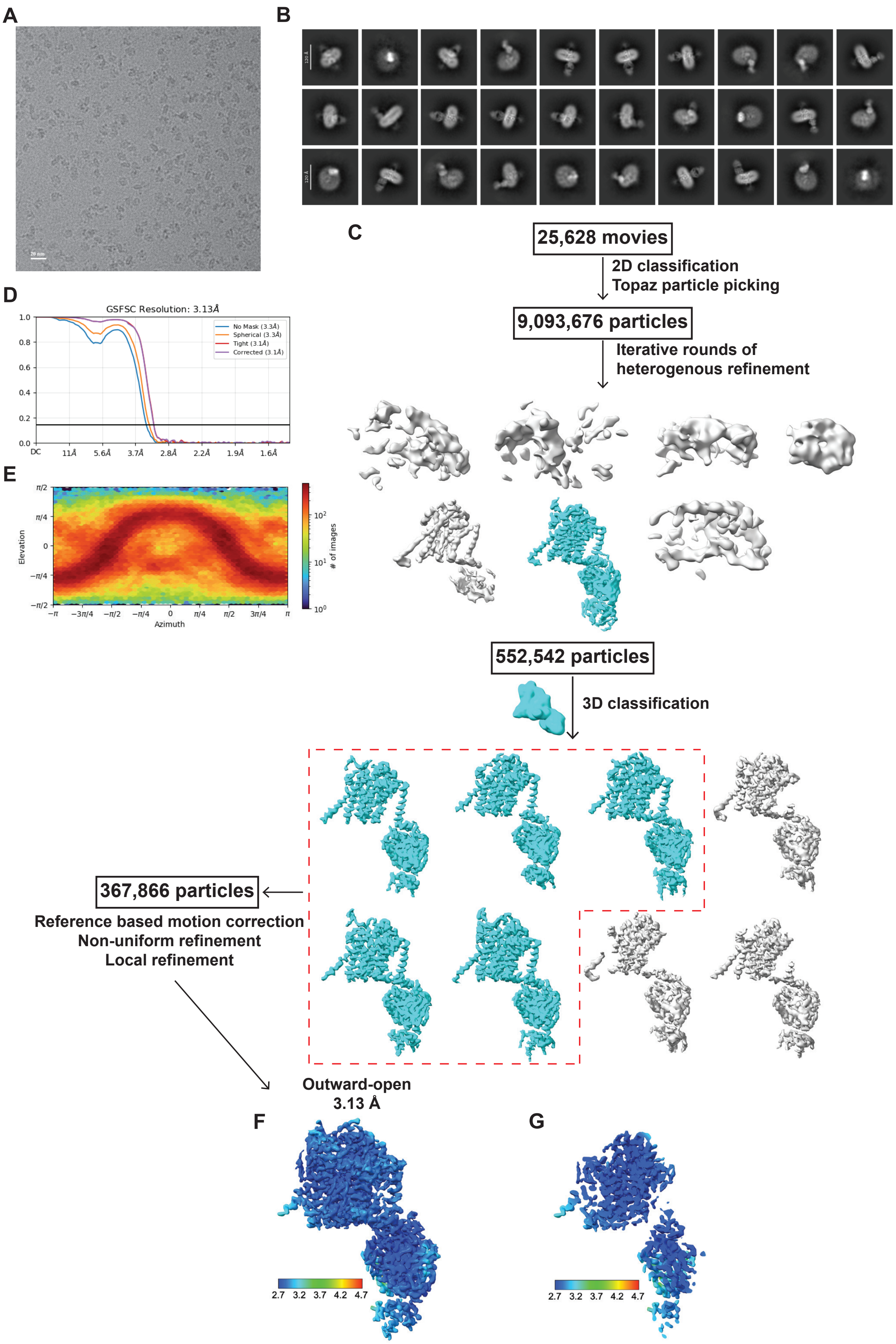
A
B
C
D
E
F
G
25,628 movies
2D classification
Topaz particle picking
9,093,676 particles
Iterative rounds of
heterogenous refinement
552,542 particles
3D classification
367,866 particles
Reference based motion correction
Non-uniform refinement
Local refinement
Outward-open
3.13 Å
GSFSC Resolution: 3.13Å
No Mask (3.3Å)
Spherical (3.3Å)
Tight (3.1Å)
Corrected (3.1Å)
Elevation
Azimuth
### of images
2.7 3.2 3.7 4.2 4.7

**Fig. S7.** Cryo-EM data processing of PFA- and $Na^{+}$-bound zfSLC34A2.

(A) Representative micrograph. (B) Representative 2D class averages. (C) Flowchart for cryo-EM data processing. (D) Gold-standard FSC curve (cutoff of 0.143) of the final density map. (E) Angular orientation distribution of all particles used in the final 3D reconstruction. (F) Final density map used for structural modelling. Surface color indicates estimated local resolution. (G) A cut-open view.

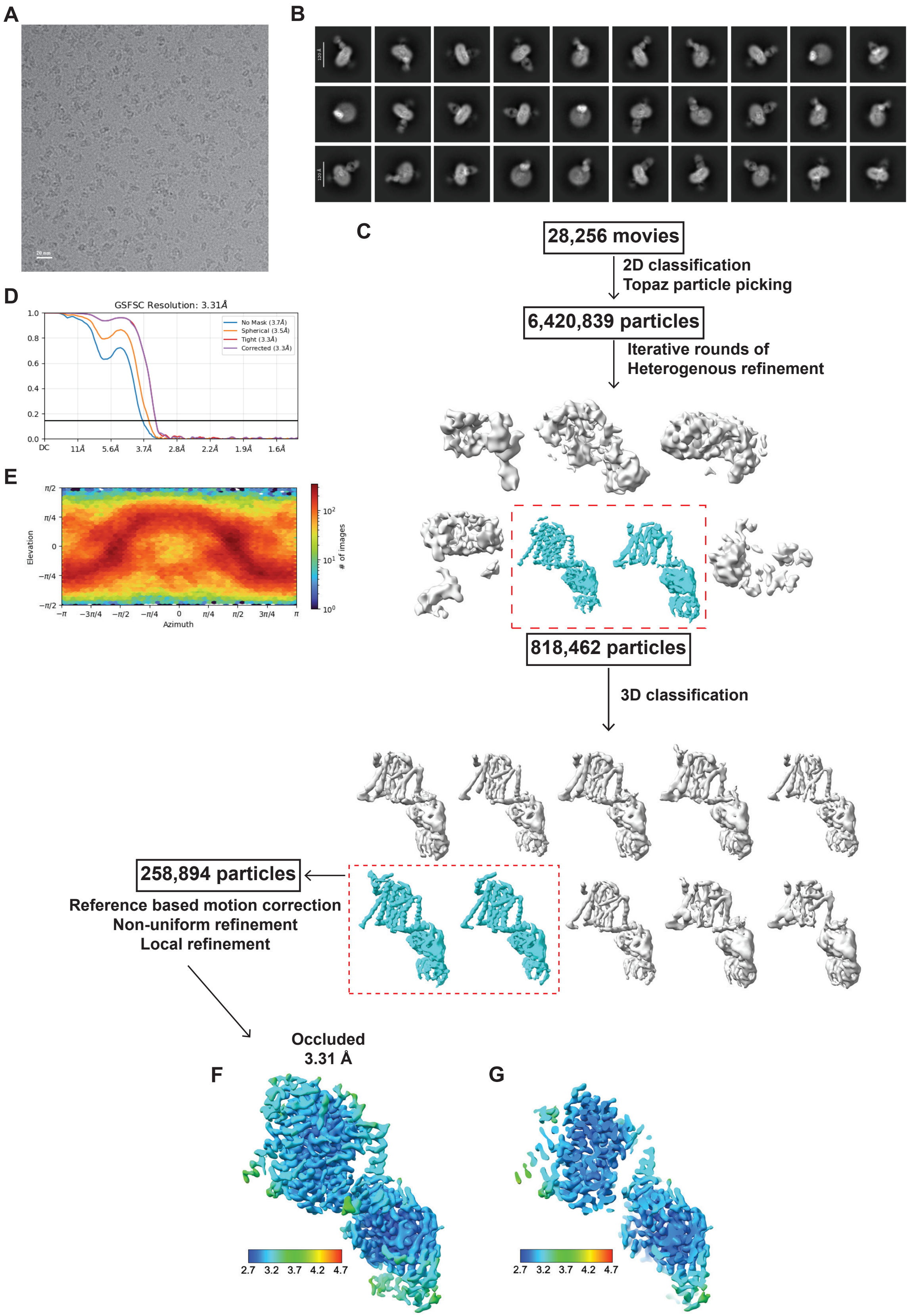
A
B
C
28,256 movies
2D classification
Topaz particle picking
6,420,839 particles
Iterative rounds of
Heterogenous refinement
818,462 particles
3D classification
258,894 particles
Reference based motion correction
Non-uniform refinement
Local refinement
D
GSFSC Resolution: 3.31Å
No Mask (3.7Å)
Spherical (3.5Å)
Tight (3.3Å)
Corrected (3.3Å)
DC
11Å
5.6Å
3.7Å
2.8Å
2.2Å
1.9Å
1.6Å
E
Elevation
Azimuth
### of images
F
Occluded
3.31 Å
2.7 3.2 3.7 4.2 4.7
G
2.7 3.2 3.7 4.2 4.7

**Fig. S8.** Cryo-EM data processing of apo zfSLC34A2 S135C.

(A) Representative micrograph. (B) Representative 2D class averages. (C) Flowchart for cryo-EM data processing. (D) Gold-standard FSC curve (cutoff of 0.143) of the final density map. (E) Angular orientation distribution of all particles used in the final 3D reconstruction. (F) Final density map used for structural modelling. Surface color indicates estimated local resolution. (G) A cut-open view.

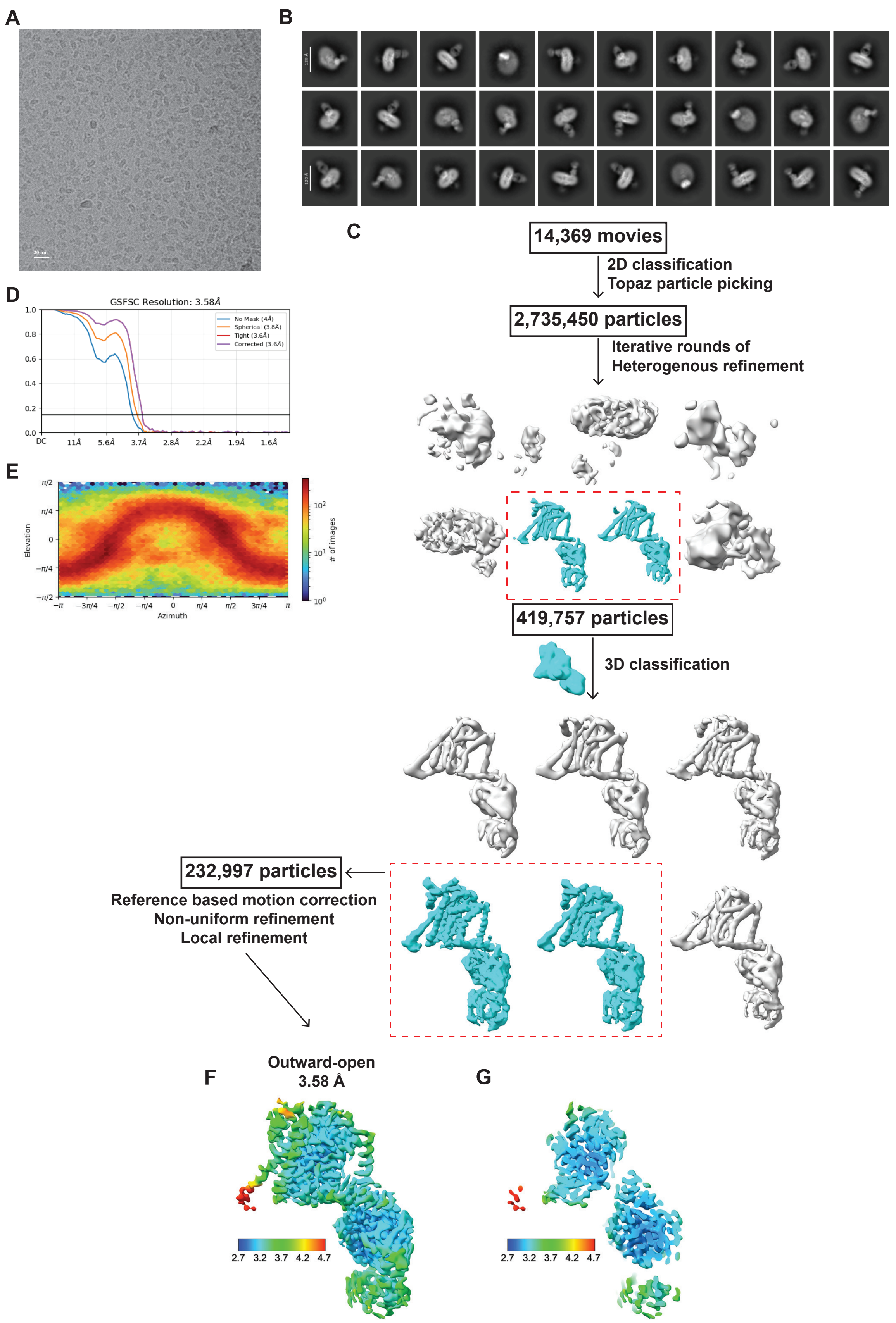
A
B
C
14,369 movies
2D classification
Topaz particle picking
2,735,450 particles
Iterative rounds of
Heterogenous refinement
419,757 particles
3D classification
232,997 particles
Reference based motion correction
Non-uniform refinement
Local refinement
D
GSFSC Resolution: 3.58Å
No Mask (4Å)
Spherical (3.8Å)
Tight (3.6Å)
Corrected (3.6Å)
E
Elevation
Azimuth
### of images
F
Outward-open
3.58 Å
2.7 3.2 3.7 4.2 4.7
G
2.7 3.2 3.7 4.2 4.7

**Fig. S9.** Cryo-EM data processing of Pi- and $Na^+$-bound zfSLC34A2 S135C.

(A) Representative micrograph. (B) Representative 2D class averages. (C) Flowchart for cryo-EM data processing. (D) Gold-standard FSC curve (cutoff of 0.143) of the final density map. (E) Angular orientation distribution of all particles used in the final 3D reconstruction. (F) Final density map used for structural modelling. Surface color indicates estimated local resolution. (G) A cut-open view.

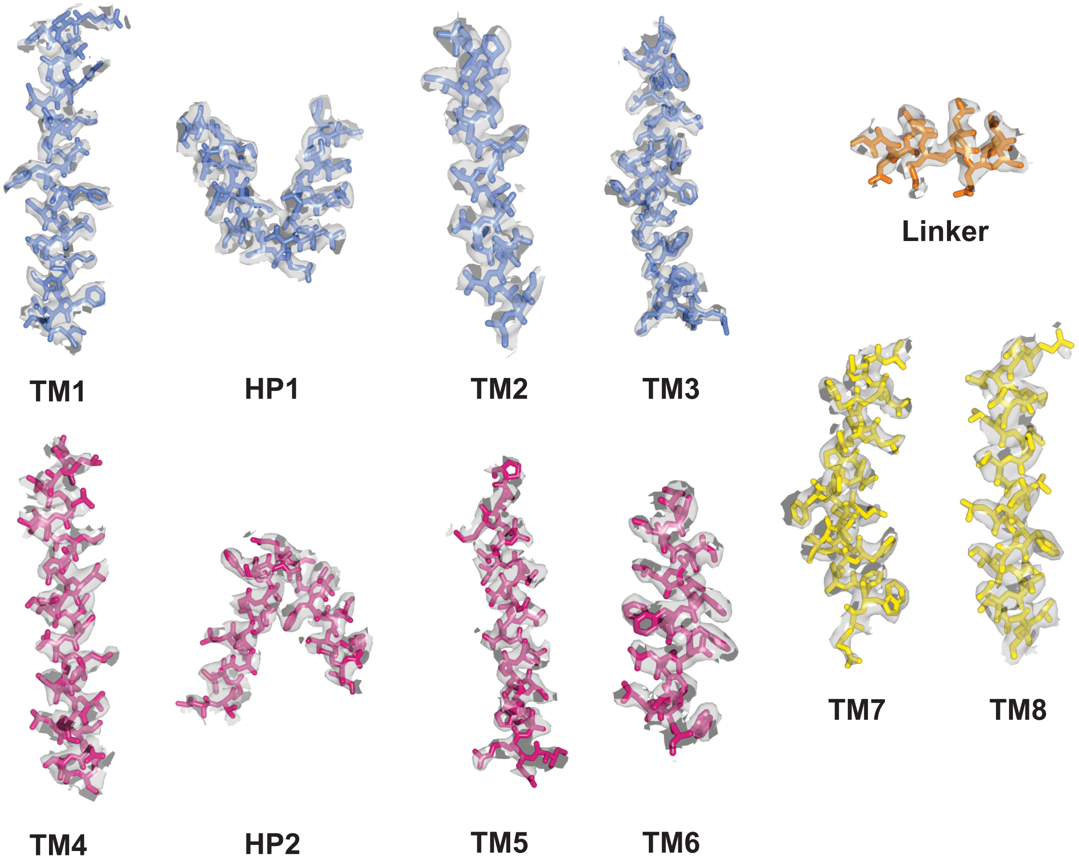


**Fig. S10.** Quality of cryo-EM density of key structural elements.

Representative segments of the cryo-EM density map from the Pi- and $Na^+$-bound zfSLC34A2 structure with the atomic model built *de novo* with assistance from ModelAngelo. Each segment is labeled and demonstrates the quality of the density-modified map (gray surface, 1$\sigma$ contour) from various regions of the reconstruction.

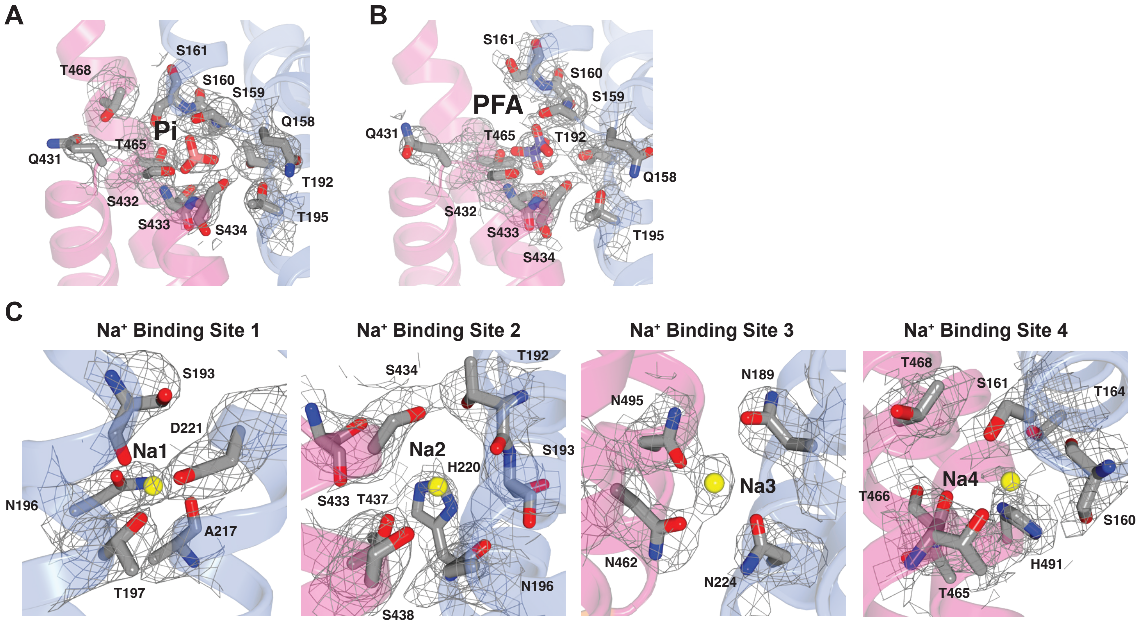


**Fig. S11.** Quality of cryo-EM density of ligands.

Representative segments of the cryo-EM density modified map (silver mesh, 1.5σ contour) from the Pi- and $Na^+$-bound zfSLC34A2 (A,C) and PFA- and $Na^+$-bound zfSLC34A2 (B) structures focused on the binding sites for the Pi, PFA, and $Na^+$.

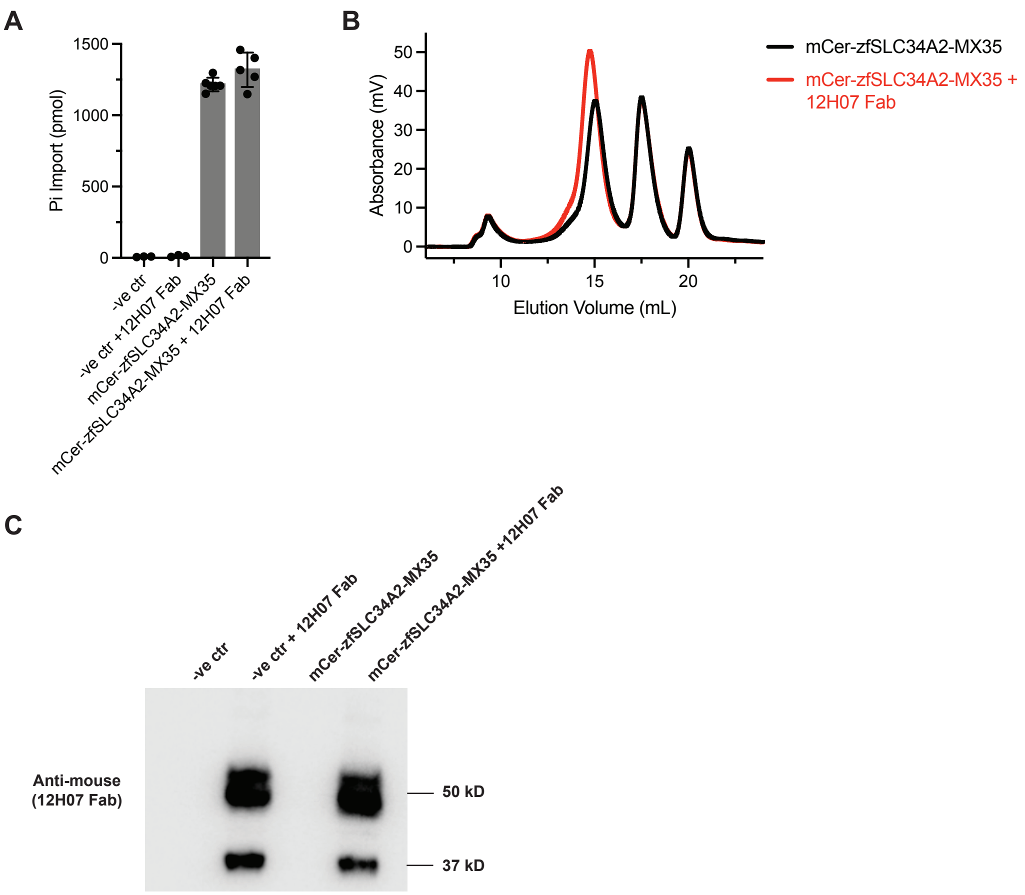


**Fig. S12.** Transport function of zfSLC34A2-MX35 is not altered in the presence of the 12H07 Fab.

(A) Pi import mediated by mCer-zfSLC34A2-MX35 in the absence and presence of the 12H07 Fab. Data are represented as the mean ± SD (n=5-7 samples). (B) FSEC analysis of samples for which function was evaluated. The proteins from three oocytes per condition were extracted in GDN. A shift in the peak indicates that the zfSLC34A2-MX35 + 12H07 Fab complex was formed. (C) Western blot analysis of the oocytes used for transport assay in A. Intact 12H07-Fab can be detected after injection.

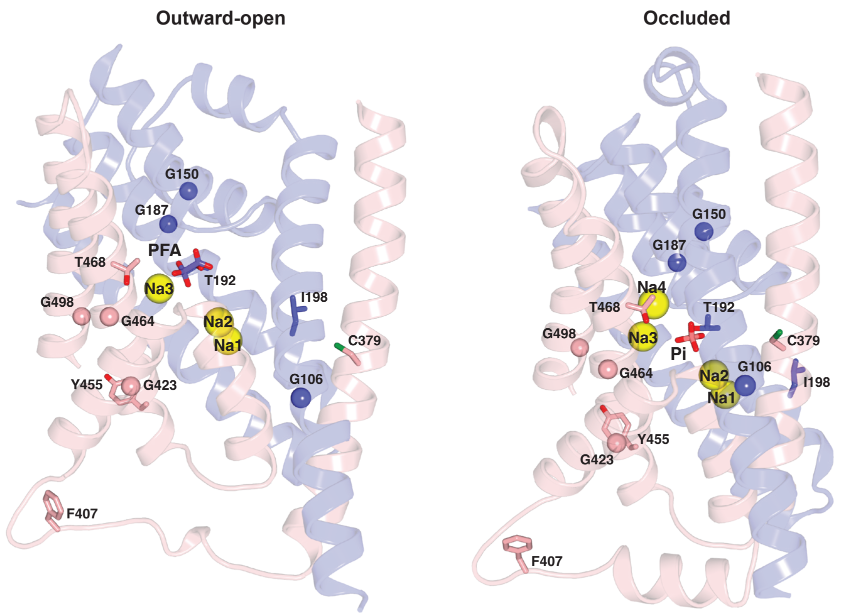


**Fig. S13.** Locations of PAM disease mutations mapped on the outward-open and occluded structures of SLC34A2.

Residues with known missense mutations that cause lung disease phenotypes are represented as sticks, or spheres in the case of glycines. All known PAM disease-associated mutations are confined to the transport domain; therefore, the extracellular and accessory domains are omitted for clarity.

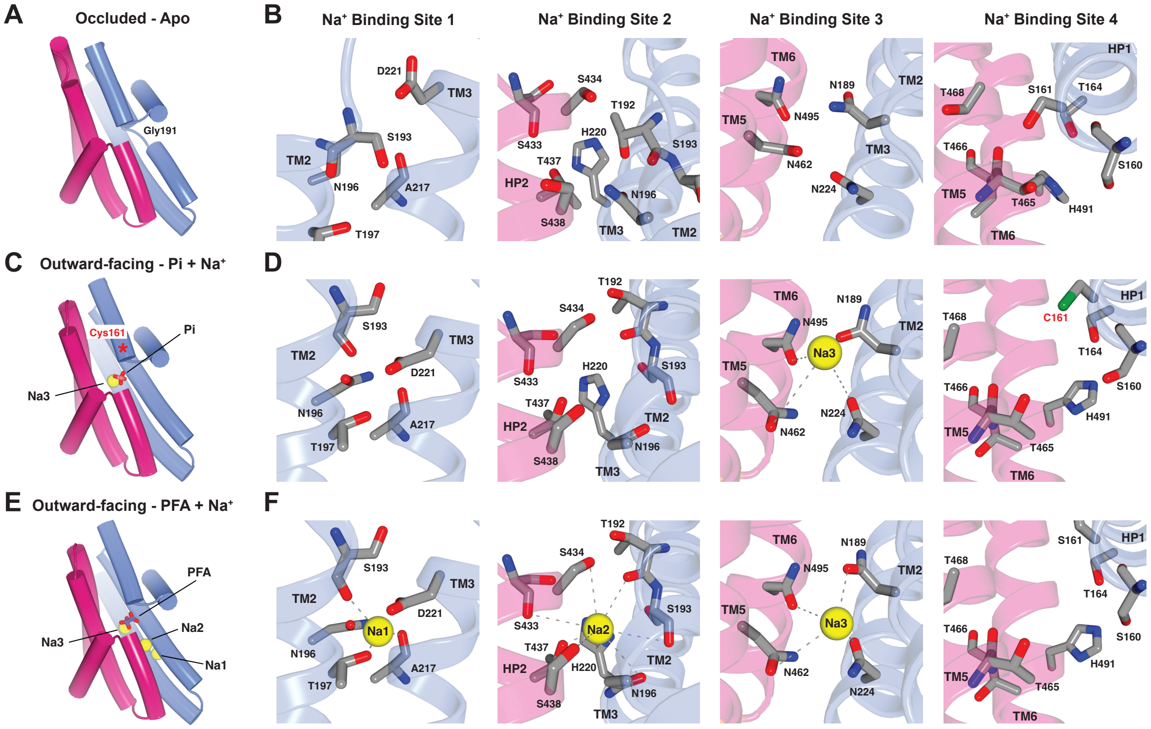


**Fig. S14. Comparison of the Na$^+$ binding sites in different structures.**

(A-F) Pi substrate (pink sticks), PFA inhibitor (purple sticks), and Na$^+$ ions (yellow spheres) binding to the transport domain of SLC34A2 in the apo (A), S135C Pi- and Na$^+$-bound (C), and PFA- and Na$^+$-bound (E) structures. Interactions with the Na$^+$ ions (yellow spheres) and the coordinating residues for Na$^+$-binding sites 1, 2, 3, and 4 are shown for all three structures, including apo (B), S135C Pi- and Na$^+$-bound (D), and PFA- and Na$^+$-bound (F). The resolution prevents direct visualization of Na$^+$ ions in binding sites 1 and 2 in the S135C Pi- and Na$^+$-bound structure. Mutation of Ser135 (Ser161 hSLC34A2 numbering) to cysteine disrupts Na4 binding.

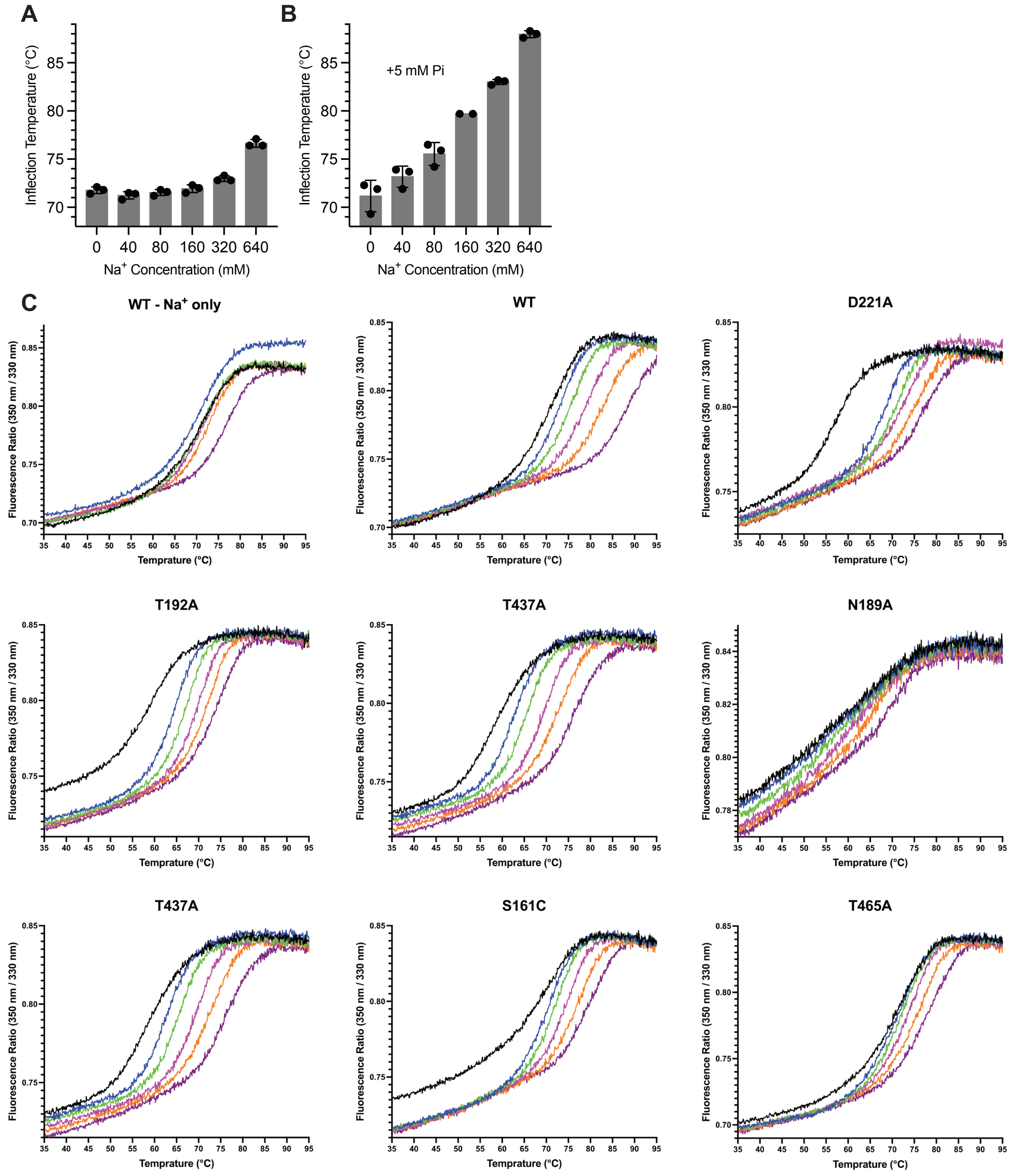


**Fig. S15.** Thermostability assays of purified hSLC34A2 to evaluate substrate and ion binding. (A) $Na^+$ dose-response curve in the absence of Pi. (B) $Na^+$ dose-response curve in the presence of 5 mM Pi. (C) Raw thermal stability assay profiles for wild-type hSLC34A2 and each mutant evaluated. Individual melting curves from nano differential scanning fluorimetry (nanoDSF) are shown. Each trace represents the change of fluorescence ratio during heating at a defined $Na^+$ concentration, colored-coded as follows: black (0 mM), blue (40 mM), green (80 mM), magenta (160 mM), orange (320 mM), and purple (640 mM).

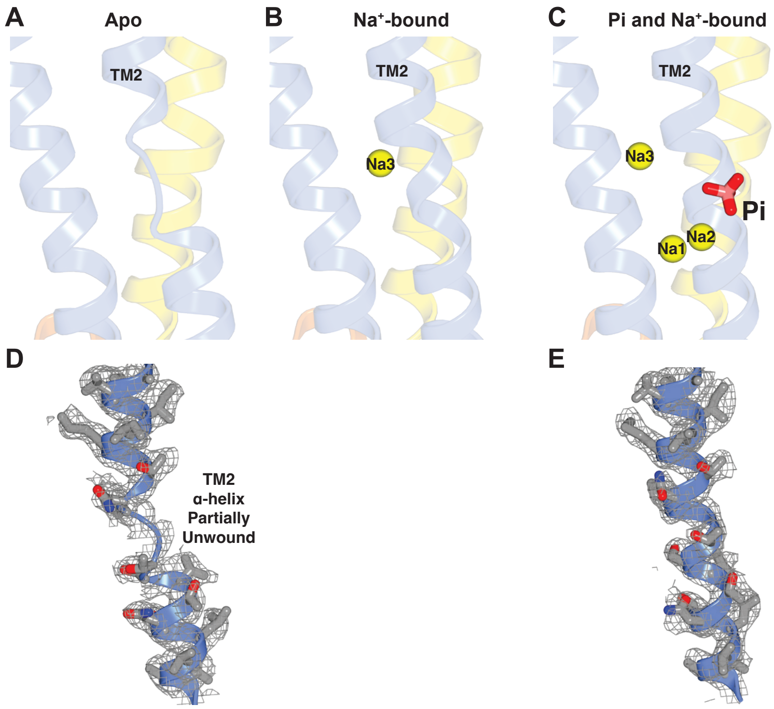


**Fig. S16.** Comparison of the apo, $Na^+$-bound, and Pi- and $Na^+$-bound structures.
(A-C) Cartoon representations of SLC34A2, highlighting the TM2 region that undergoes local unwinding, in the apo (A), $Na^+$-bound (B), and Pi- and $Na^+$-bound samples (C). Although the moderate resolution precludes direct visualization of $Na^+$ ions in binding sites 1 and 2 in the $Na^+$-bound structure, the observed conformational change, specifically, the continuity of TM2 as an that $\alpha$-helix, suggests that ion binding, rather than Pi binding, drives this structural rearrangement. (D-E) Representative segments of the cryo-EM density modified map (silver mesh, 1.5$\sigma$ contour) from the apo zfSLC34A2 (D) and Pi- and $Na^+$-bound zfSLC34A2 (E) structures, focusing on TM2, which transitions from an unwound conformation to an $\alpha$-helix upon $Na^+$ ion binding.

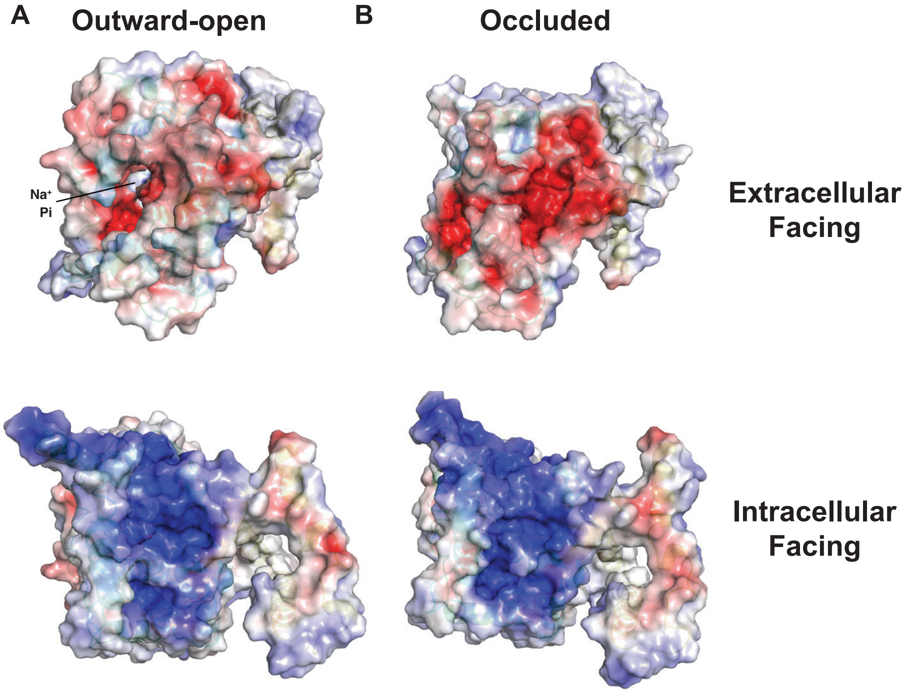


**Fig. S17.** Electrostatics of the entrance and exit of the substrate translocation pathway.
In both the outward-facing (A) and occluded states (B), the extracellular face of SLC34A2 is negatively charged and the intracellular face is positively charged. The molecular surface is colored according to electrostatic potential: light gray regions are neutral; red, -5 $kTe^{-1}$; blue, +5 $kTe^{-1}$.

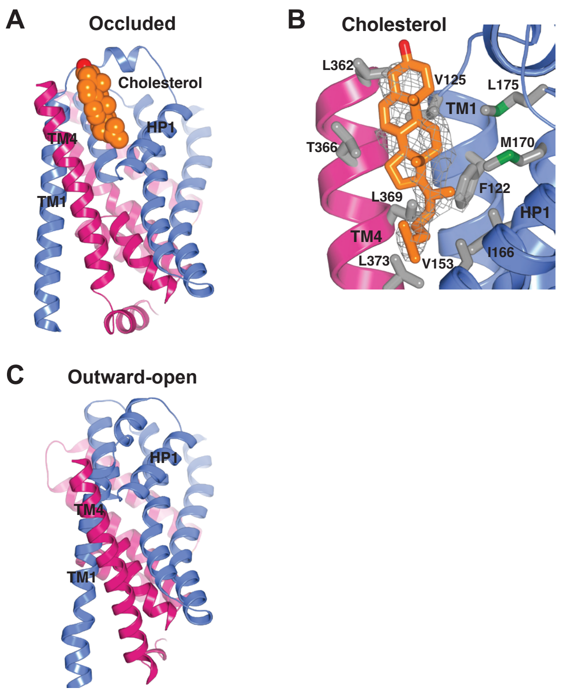


**Fig. S18.** Cholesterol-protein interaction specific to the occluded state.

(A) A cholesterol molecule (orange) packs between TM1, HP1, and TM4 in the occluded state. (B) The cholesterol molecule is situated in a cavity lined with hydrophobic residues. Cholesterol density is shown (gray mesh, 1.3σ contour). (C) This cholesterol-binding cavity is absent in the outward-open state.

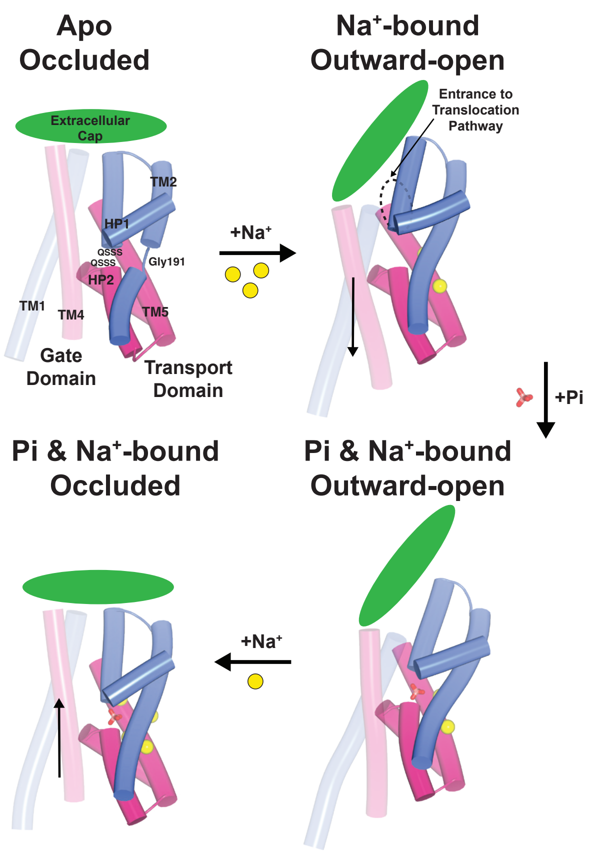


**Fig. S19.** Mechanism of substrate and ion loading, coupling, and conformational changes associated with SLC34A2 transport.

Binding of Na+ ions to sites 1, 2, and 3, induces the transition of an unwound segment in TM2, centered on Gly191, to $\alpha$-helical. This structural change creates a favorable binding environment for Pi at the junction of the helical hairpins, which is defined by the QSSS repeat motifs. Finally, the binding of a fourth $Na^+$ ion facilitates the transition from the outward-open to the occluded state by narrowing the entrance pathway between HP1 and TM5. This transition is accompanied by a significant repositioning of TMs 1 and 4, as well as the extracellular cap to further restrict substrate and ion access.

**Table S1.** Cryo-EM data collection, refinement, and validation statistics.

| | #1 SLC34A2<br><br>Occluded<br>(EMDB-73829)<br>(PDB 9Z60) | #2 SLC34A2<br>+Na$^+$<br>Outward-open<br>(EMDB-73830)<br>(PDB 9Z61) | #3 SLC34A2<br>+Pi +Na$^+$<br>Occluded<br>(EMDB-73831)<br>(PDB 9Z62) | #4 SLC34A2<br>+PFA +Na$^+$<br>Outward-open<br>(EMDB-73832)<br>(PDB-9Z63) |
|---|---|---|---|---|
| **Data collection and processing** | | | | |
| Magnification | 105,000 | 165,000 | 165,000 | 165,000 |
| Voltage (kV) | 300 | 300 | 300 | 300 |
| Electron exposure (e–/Å$^2$) | 53 | 60 | 60 | 60 |
| Defocus range (μm) | -0.7 to -1.7 | -0.7 to -1.7 | -0.7 to -1.7 | -0.7 to -1.7 |
| Pixel size (Å) | 0.826 | 0.725 | 0.725 | 0.725 |
| Symmetry imposed | C1 | C1 | C1 | C1 |
| Initial particle images (no.) | 5,395,965 | 4,689,640 | 3,963,529 | 9,093,676 |
| Final particle images (no.) | 139,524 | 336,681 | 556,732 | 367,866 |
| Map resolution (Å)<br>FSC threshold 0.143 | 3.53 | 3.68 | 3.10 | 3.02 |
| Map resolution range (Å) | 3.0-6.3 | 3.1-7.5 | 2.6-7.5 | 2.6-3.8 |
| **Refinement** | | | | |
| Initial model used (PDB code) | ModelAngelo | ModelAngelo | ModelAngelo | ModelAngelo |
| Model resolution (Å)<br>FSC threshold 0.5 | 3.63 | 3.86 | 3.20 | 3.07 |
| Map sharpening *B* factor (Å$^2$) | -140.4 | -142.3 | -105.4 | -114.1 |
| Model composition | | | | |
| Non-hydrogen atoms | 5326 | 5388 | 5359 | 5397 |
| Protein residues | 694 | 705 | 696 | 705 |
| Ligands | 1 | 1 | 6 | 4 |
| *B* factors (Å$^2$) | | | | |
| Protein | 52.81 | 69.58 | 52.37 | 41.52 |
| Ligand | 57.62 | 70.34 | 62.24 | 28.67 |
| R.m.s. deviations | | | | |
| Bond lengths (Å) | 0.002 | 0.002 | 0.002 | 0.003 |
| Bond angles (°) | 0.486 | 0.461 | 0.470 | 0.480 |
| Validation | | | | |
| MolProbity score | 1.31 | 1.49 | 1.35 | 1.44 |
| Clashscore | 5.75 | 6.06 | 4.89 | 5.96 |
| Poor rotamers (%) | 0.52 | 0.68 | 0.68 | 0.68 |
| Ramachandran plot | | | | |
| Favored (%) | 98.10 | 97.13 | 97.52 | 97.42 |
| Allowed (%) | 1.90 | 2.87 | 2.48 | 2.58 |
| Disallowed (%) | 0.00 | 0.00 | 0.00 | 0.00 |

| | #5 SLC34A2 S135C Occluded (EMDB-73833) (PDB 9Z64) | #6 SLC34A2 S135C +Pi +Na$^+$ Outward-open (EMDB-73833) (PDB 9Z66) |
|---|---|---|
| **Data collection and processing** | | |
| Magnification | 165,000 | 165,000 |
| Voltage (kV) | 300 | 300 |
| Electron exposure (e–/Å$^2$) | 61.19 | 59.81 |
| Defocus range (μm) | -0.7 to -1.7 | -0.7 to -1.7 |
| Pixel size (Å) | 0.725 | 0.725 |
| Symmetry imposed | C1 | C1 |
| Initial particle images (no.) | 6,420,839 | 2,735,450 |
| Final particle images (no.) | 258,894 | 232,997 |
| Map resolution (Å) FSC threshold 0.143 | 3.11 | 3.31 |
| Map resolution range (Å) | 2.7-5.4 | 2.8-6.0 |
| **Refinement** | | |
| Initial model used (PDB code) | ModelAngelo | ModelAngelo |
| Model resolution (Å) FSC threshold 0.5 | 3.27 | 3.43 |
| Map sharpening *B* factor (Å$^2$) | -121.0 | -121.2 |
| Model composition | | |
| Non-hydrogen atoms | 5337 | 5393 |
| Protein residues | 694 | 705 |
| Ligands | 1 | 2 |
| *B* factors (Å$^2$) | | |
| Protein | 47.65 | 71.57 |
| Ligand | 54.15 | 56.55 |
| R.m.s. deviations | | |
| Bond lengths (Å) | 0.002 | 0.002 |
| Bond angles (°) | 0.449 | 0.431 |
| Validation | | |
| MolProbity score | 1.23 | 1.53 |
| Clashscore | 4.35 | 7.07 |
| Poor rotamers (%) | 0.51 | 0.68 |
| Ramachandran plot | | |
| Favored (%) | 97.95 | 97.27 |
| Allowed (%) | 2.05 | 2.73 |
| Disallowed (%) | 0.00 | 0.00 |
